# A red-emitting, genetically encoded indicator for two-photon voltage recording in vivo

**DOI:** 10.64898/2026.06.01.726307

**Authors:** Shuyuan Yang, Alex James McDonald, Xiaoyu Lu, Vincent Villette, Noura Hakam, Mario Galdamez, Tanmayee Torne-Srivas, Gregory Foran, Xiaoyu Dong, Matthew Shorey, Nima Bavili, Shujuan Lai, Zhuohe Liu, Haixin Liu, Tom Maslianitsyna, Rémi Fournel, Emiliano Ronzitti, Jun Zhu, Ryan Gregory Natan, Jonathan Bradley, Daniela Gaspar Santos, Yu-Yau York Shan, Lindsey Ran, Ming Hu, Valentina Emiliani, Na Ji, Jacob Reimer, Laurent Bourdieu, François St-Pierre

## Abstract

Genetically encoded voltage indicators (GEVIs) enable minimally invasive, cell-type-specific optical measurements of neuronal membrane potential with millisecond temporal resolution. Red-shifted GEVIs are especially advantageous because they permit spectral multiplexing with complementary sensors and enable all-optical circuit interrogation in combination with blue-light-activated opsins. Despite these advantages, existing red GEVIs remain poorly suited for in vivo use due to limited performance under two-photon (2P) excitation, the predominant modality for deep-tissue imaging. Here, we introduce VADER1, a red GEVI that overcomes this limitation and enables reliable spike detection in vivo under 2P illumination. Under 2P excitation, VADER1 supports extended voltage imaging with both random-access and resonant-scanning microscopy, enables recordings from neurons as deep as cortical layer 5, and allows dual-color imaging with calcium indicators. By filling a critical spectral gap, VADER1 enables integrated optical measurements of fast electrical activity alongside other neural signals and establishes a foundation for two-photon all-optical electrophysiology.

## Introduction

Genetically encoded voltage indicators (GEVIs) are transforming neuroscience by enabling minimally invasive, cell-type-specific optical recordings of membrane-potential dynamics with high temporal fidelity and subcellular resolution^1,2^. Although GEVI development efforts have emphasized green indicators^3–5^, red GEVIs would enable spectral multiplexing with green GEVIs to report from different cell types or compartments (e.g., axons versus dendrites) simultaneously. Red GEVIs could also be paired with green reporters of other modalities when suitable red-shifted alternatives are lacking, and are optimally suited for all-optical experiments that combine voltage imaging with optogenetic control of neural activity. Because even red-shifted opsins absorb appreciably in the blue, pairing them with green GEVIs can introduce significant optical cross-talk^6^, compromising artifact-free voltage —such as those needed for in vivo circuit mapping. By contrast, combining red GEVIs with blue-absorbing opsins, which show minimal activation by green light^7^, offers a cleaner spectral separation and is often the preferred combination ^8–10^.

Despite these advantages, there are still no genetically encoded red indicators that broadly support robust voltage imaging in vivo under two-photon (2P) excitation, the dominant modality for deep-tissue functional imaging. Early attempts to engineer red GEVIs by incorporating voltage-sensing domains from voltage-sensitive phosphatases produced several sensors whose modest responsiveness did not warrant in vivo benchmarking^11–13^. In contrast, rhodopsin-based GEVIs have yielded several red indicators—such as the Arch series^14–20^, VARNAM^13,21^, Cepheid1b/1s^22^, and pAceR²³—that exhibit robust one-photon voltage responses in vivo. However, rhodopsin-based sensors exhibit minimal voltage sensitivity under standard 2P illumination methods^23,24^. Although recent studies have demonstrated that specialized 2P imaging modalities can partially rescue their lower 2P performance^25^, these approaches often require non-standard optical systems, the resulting voltage responses remain small, and some designs rely on exogenous dyes^26^. Moreover, rhodopsin-based GEVIs display complex photophysics under 2P excitation, presenting response amplitudes that vary with illumination intensity^27^. Together, these limitations hinder the practical deployment of red GEVIs for routine in vivo experiments by the broader neuroscience community.

Here, we report the development of VADER1, the first fully genetically encoded red GEVI capable of robust, long-duration 2P voltage imaging in vivo. VADER1 was evolved using a custom high-throughput screening platform that simultaneously evaluates brightness, kinetics, and the fluorescence–voltage relationship, enabling the identification of variants with superior optical and biophysical properties. VADER1 produces spike-evoked fluorescence changes with a signal-to-noise ratio comparable to that of the established green GEVI JEDI-2P^28^, and supports stable voltage recordings for up to one hour using ultrafast local volume excitation (ULoVE) microscopy. In addition, VADER1 reliably reports spikes from deep cortical layers under conventional resonant-scanning 2P imaging. We further demonstrate its suitability for multiplexed experiments by combining VADER1 with the green calcium indicator GCaMP6s. Together, these results show that VADER1 fills a critical gap in the 2P voltage-imaging toolkit and provides a foundation for in vivo, spectrally multiplexed functional 2P optical recordings.

## Results

### High-throughput multiparametric optimization of a red, positive-going voltage spike detector

We hypothesized that a general-use, red-shifted GEVI suitable for 2P microscopy could be engineered using the JEDI/ASAP indicator architecture, which inserts a circularly permuted fluorescent protein into a short extracellular loop of a voltage-sensing domain (VSD). Compared with *A. victoria*-based GFP, orange and red fluorescent proteins (OFPs/RFPs) are often more prone to aggregation or oligomerization and less tolerant to fusion^29^. We thus first screened voltage-sensing domains incorporating OFPs and RFPs^30–35^ for efficient plasma membrane targeting (Figure S1.1A). As anticipated, several candidates—including TagRFP-T and mCherry—generated substantial unwanted intracellular fluorescence. Of the 10 fluorescent proteins tested, fusions based on mApple and mOrange exhibited the most elicient plasma membrane localization, comparable to a standalone RFP fused to a membrane-anchoring prenylation motif (Figure S1.1B).

We selected mApple for red GEVI development due to its more red-shifted spectra, which facilitates multiplexing, and because it has proven amenable to engineering in other biosensor applications^36–41^. We replaced the cpGFP in JEDI-2P with a circularly permuted mApple variant from the dopamine sensor rGRAB_DA1m_^41^ (Figures 1A, S1.1C). The resulting indicator was dim and produced a small (∼10% ΔF/F_0_) dim-to-bright response to a 20-ms electric field stimulation (EFS) pulse (**Error! Reference source not found.**.1D). We designated this prototype red GEVI Voltage-Activated Domain Emitting Red 0.1 (VADER0.1).

**Figure 1.**
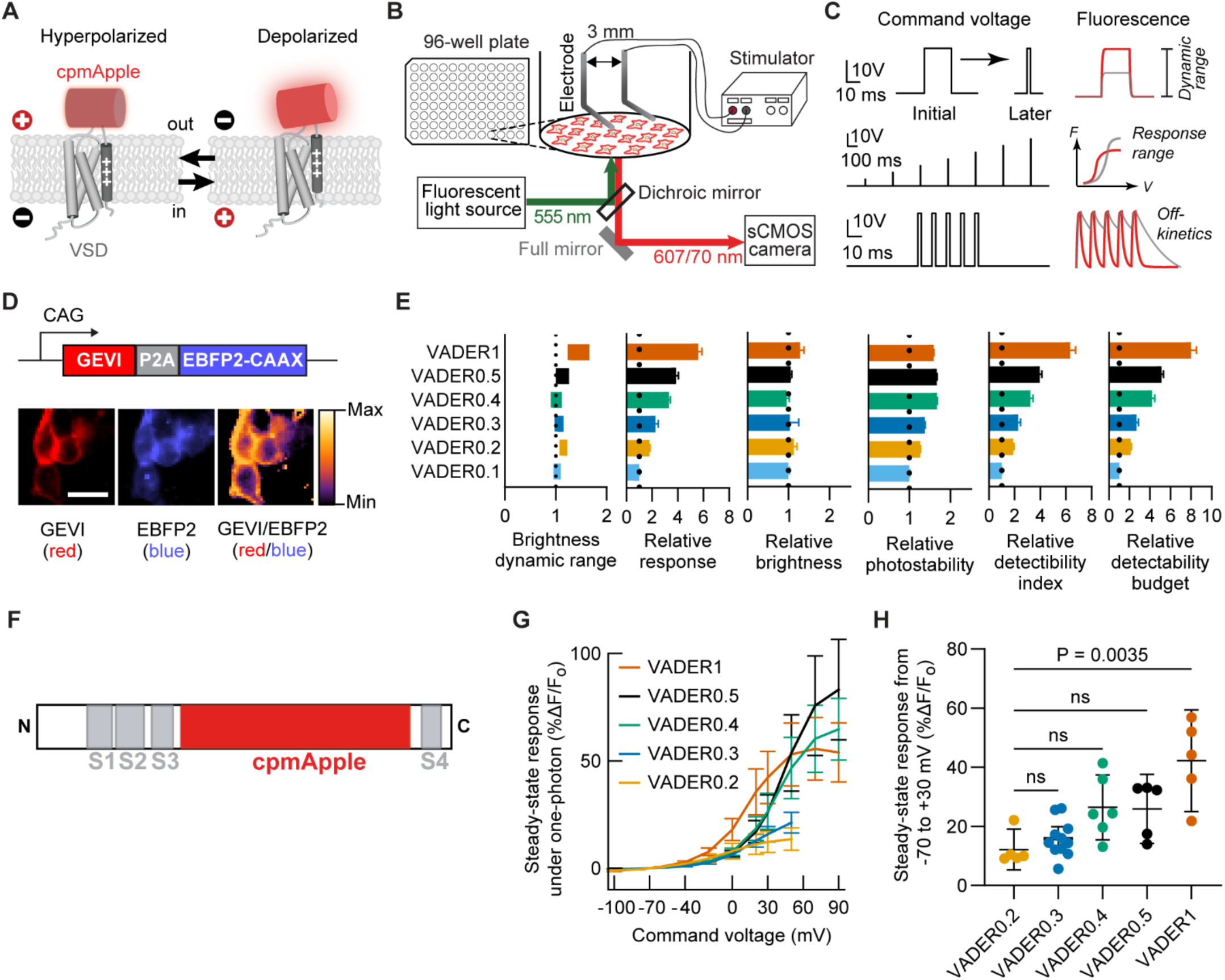
Direct evolution of a red-emitting GEVI under one-photon. (A) Conceptual schematic of the red GEVIs design used in this study. Voltage-dependent conformational changes in the voltage-sensing domain (VSD) modulate mApple fluorescence. (B) Screening setup. Fluorescence from GEVI-expressing cells was acquired by camera while membrane voltage was perturbed via electrical field stimulation (EFS). Imaging and stimulation were fully automated. (C) EFS paradigms used during screening. *Top*, single, high-amplitude pulses to assess maximal responses. Pulse duration was decreased after GEVI performance increased. *Middle,* voltage ramp to evaluate shifts in the voltage-response range. *Bottom,* pulse train used to discard variants with excessively slow kinetics. (D) Brightness quantification strategy. *Top*, screening cassette schematic. Red GEVIs were co-expressed with a membrane-anchored blue fluorescent protein (EBFP2-CAAX) via a ribosome-skipping sequence (P2A). *Bottom*, Brightness was quantified as the ratio of red to blue fluorescence to reduce cell-to-cell expression variability. Scale bar: 10 μm. (E) Comparison of VADER1 and intermediate variants in the screening assay, normalized to VADER0.1. Brightness dynamic range: bars indicate GEVI brightness (B) from the dim (left edge) to the bright (right edge) states, normalized to VADER0.1’s dim state. Response: Peak Δ𝐹/𝐹_0_ evoked by a 2.5-ms, 15-V EFS pulse. Photostability (𝑃): area under the normalized baseline brightness curve during the 9-s EFS assay. Detectability index^42,43^: Δ𝐹/𝐹_0_ · √𝐵. Detectability budget: 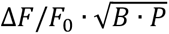. n = 5 (VADER0.2) or 6 (other variants) independently transfected wells. Error bars: 95% CI. (F) Linear protein schematic of the VADER family of indicators. S1-S4: transmembrane segments of the voltage-sensing domain. (**G**-**H**) Characterization by whole-cell voltage clamp combined with 1P imaging at ∼22°C. n = 5-11 HEK293A cells. Error bars: 95% CI. (G) Mean steady-state fluorescence response to 1-s voltage steps from -70 mV. (H) Quantification of (G) for 100-mV voltage steps. Black lines: median. ns, non significant (P > 0.05; Kruskal–Wallis test followed by Dunn’s multiple comparisons test).

We optimized VADER0.1 performance by screening single- and double-site high-throughput saturation-mutagenesis libraries in 96-well plates, simultaneously evaluating brightness, photostability, sensitivity, voltage range, and kinetics. Screening was performed under one-photon widefield excitation because the laser in our 2P laser-scanning platform^42^ provided insufficient power at RFP-excitation wavelengths. We adapted the one-photon system previously developed for screening the green indicator JEDI-1P ^43^, modifying illumination and filter wavelengths for red indicators (Figure 1B).

As the directed evolution campaign improved indicator performance, we implemented more stringent assays, progressing from long (20-ms) to short (2.5-ms) EFS pulses (Figure 1C, top). For libraries screened later in VADER development, we introduced stimulation protocols to detect shifts in the voltage-response range (Figure 1C, middle) and to identify variants with excessively slow kinetics (Figure 1C, bottom). To normalize brightness measurements across cells and wells, VADER variants were co-expressed with a membrane-anchored blue fluorescent protein (Figure 1D). Photostability was quantified by measuring fluorescence decay during the 8-s exposure screening trials.

We screened 53 libraries, targeting 29 residues in the VSD and 13 residues in cpmApple. Indicator performance improved across multiple rounds of optimization (Figure 1E). We identified six beneficial modifications around the junctions between the VSD and cpmApple (Figure 1F). Although shortening linker sequences between sensing and optical domains often enhance performance^44^, both VSD-cpmApple junctions benefited from the insertion of one additional residue. We discovered three advantageous mutations targeting the RFP domain.

Voltage-response curves of VADER0.1–0.4 remained right-shifted relative to the physiological range (Figure 1G). Therefore, we designed libraries targeting the N- and C-terminal regions of the VSD fourth transmembrane helix (S4) to identify mutations promoting its activated (Up) state at lower voltages. One of these mutations, located at the C-terminal end of S4, produced a pronounced leftward shift. This mutation yielded strong sensitivity within the suprathreshold voltage range (Figure 1G, H. Figure S1.2). We named this final, spike-optimized variant VADER1.

### VADER1 reports spike waveforms under 2P resonant scanning in vitro

We evaluated VADER1 under resonant-scanning 2P excitation and benchmarked it against FlicR2^13^ (Figure 2A), the best-performing red GEVI for 2P resonant-scanning voltage imaging. To determine optimal illumination wavelengths, we measured the 2P excitation spectrum for each indicator (Figure 2B). VADER1’s profile closely matched mApple’s, with a modest blue shift, consistent with their 1P spectra (Figure S2.1). Exciting VADER1 between 1040 and 1130 nm produced fluorescence exceeding 80% of the maximal brightness (Figure 2B). VADER1’s spectrum displayed prominent peaks at both 1055 and 1120 nm, whereas FlicR2’s spectrum was dominated by a single peak near 1075 nm. Emission spectra were largely similar among the three proteins, although FlicR2 showed a slight red shift.

**Figure 2.**
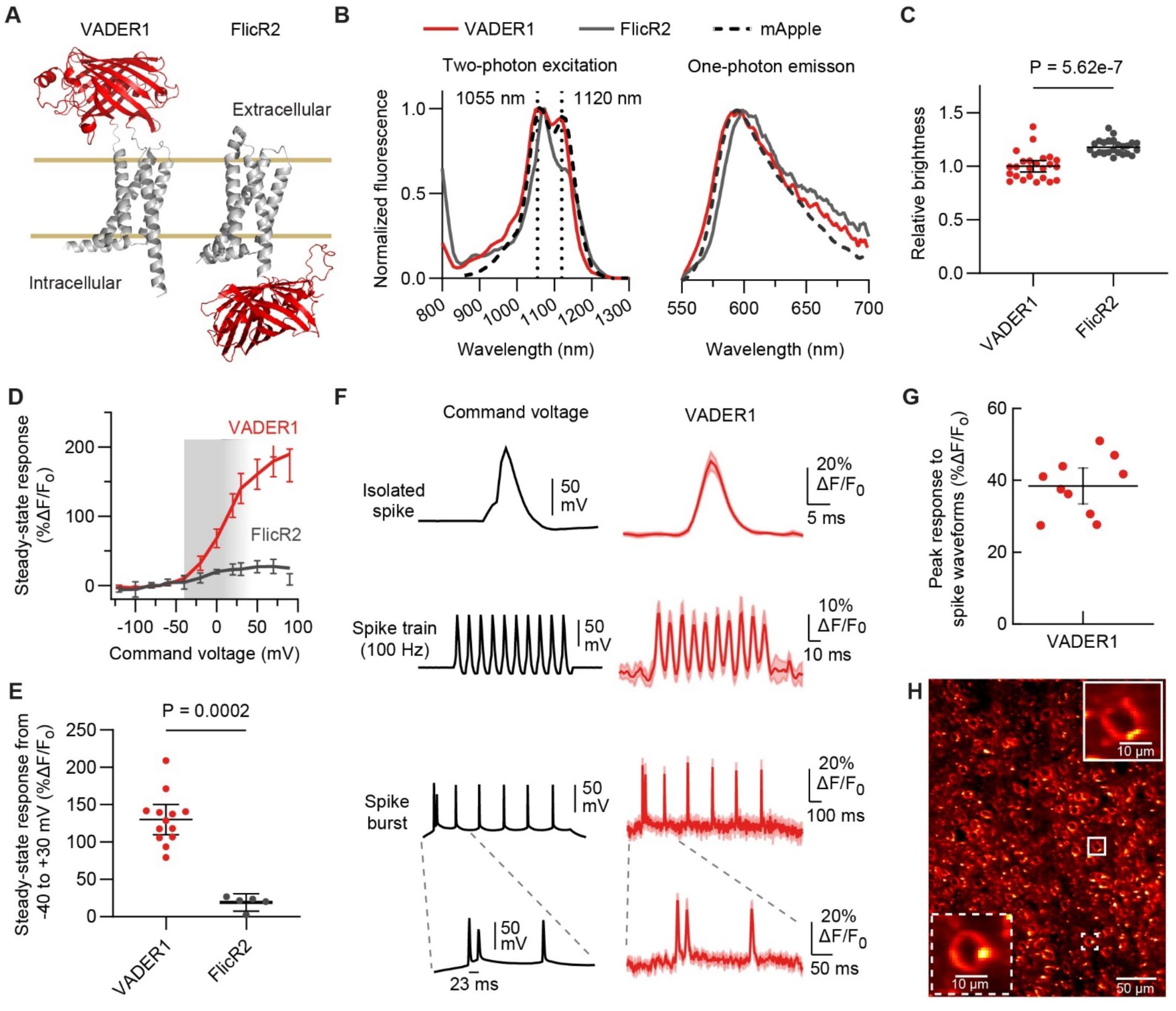
Two-photon characterization of VADER1. All in vitro data were collected at ∼22°C. (A) AlphaFold3-predicted structures. Brown lines indicate approximate plasma membrane boundaries. (B) *Left*: 2P excitation spectra. n = 45 (VADER1), 33 (FlicR2), 31 (mApple) CHO cells. *Right*: Emission spectra collected with 525/10-nm excitation. n = 6 (VADER1), 6 (FlicR2), 3 (mApple) independently transfected wells with HEK293A cells. (C) Comparison of VADER1 and FlicR2 brightness at −70 mV, corrected for differences in relative excitation efficiency at the 1050-nm illumination wavelength. Data points: means from >100 HEK-Kir2.1 cells per independently transfected well. Black lines: means across n = 24 wells. Error bars: 95% CI. P value from Welch’s t-test on log-transformed values (**D-G**) Whole-cell voltage-clamp measurements combined with simultaneous 2P imaging at 1,050 nm in mammalian cells. (D) Steady-state fluorescence–voltage relationships measured during 1-s voltage steps from −70 mV. Gray shaded region: target voltage range. n = 13 (VADER1) and 5 (FlicR2) cells. (E) Quantification of (D) for 70-mV voltage steps. Black lines: median. Error bars: 95% CI. P value from two-tailed Mann–Whitney U test (U = 0). (F) Responses to spike waveforms while imaging by resonant scanning at 440 Hz. Top row: isolated spikes (100 mV amplitude, 4-ms full width at half maximum, FWHM). Second row: 100-Hz spike train (2-ms FWHM). Third row: burst firing superimposed on subthreshold depolarization (3–4-ms FWHM). Bottom row: expanded view of the burst shown above. Dark traces indicate means; shaded regions indicate 95% CI. n = 10 VADER1 cells. (G) Quantification of peak response amplitude to isolated spikes shown in (F). Error bars: 95% CI. (H) Representative 2P image of VADER1-tKv-expressing neurons in a head-fixed mouse, following AAV injection in Layer 2/3 of visual cortex area V1. Excitation: 1035 nm. 2 additional images are shown in Fig. S2.4.

Basal brightness under 2P was measured at –70 mV to approximate the resting membrane potential of cortical pyramidal neurons. VADER1 and FlicR2 had comparable 2P brightness at their respective peak excitation wavelengths (Figure 2C). Voltage sensitivity was then assessed under whole-cell voltage clamp during 2P illumination at 1050 nm. From -120 to +90 mV, VADER1 produced a 192 ± 25% ΔF/F response (mean ± 95% CI), with most of its dynamic range (130 ± 20 % ΔF/F, mean ± 95% CI) occurring within the suprathreshold voltage window (–40 to +30 mV), underscoring its utility for spike detection. In contrast, FlicR2 exhibited only 19 ± 11% ΔF/F (mean ± 95% CI) across the same suprathreshold voltage range, ∼7.5-times lower than VADER1 (Figure 2D, E).

Because submillisecond-resolution 2P kinetic measurement techniques are not yet established, we assessed indicator kinetics under widefield one-photon illumination. FlicR2’s voltage responses were too small relative to baseline noise to permit reliable time-constant estimation. In contrast, VADER1 displayed well-resolved kinetics: a monoexponential activation time constant of ∼1.4 ms for a 100-mV depolarization step at 33 °C, and biexponential repolarization kinetics dominated by a rapid ∼0.8-ms component (Table 1). These kinetics are suitable for reporting action-potential, although kilohertz sampling rates will be required to capture maximal ΔF/F amplitudes.

**Table 1.**
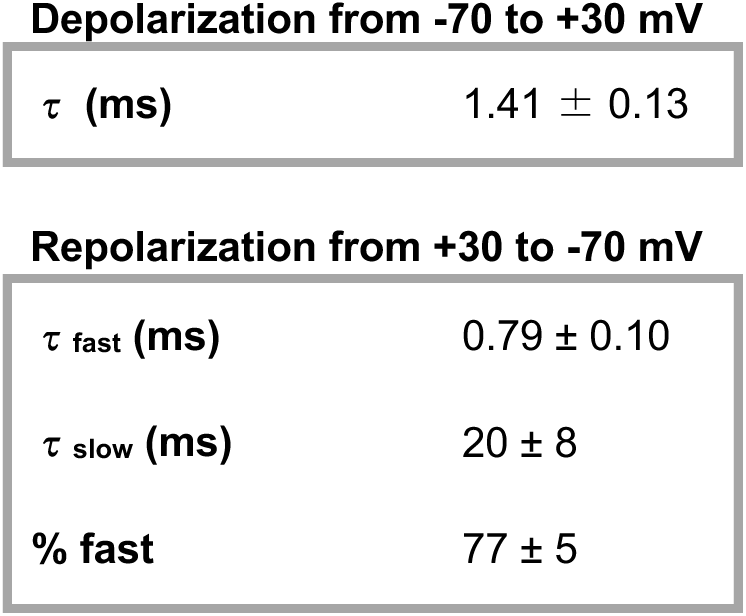
VADER1’s kinetics at 33°C under widefield 1P illumination. Voltage was modulated under whole-cell voltage clamp. Depolarization was best fit by a monoexponential function, whereas repolarization was best described by a biexponential function. Values are the mean ± 95% CI. n = 13 HEK293A cells.

**Depolarization from -70 to +30 mV**
|  |  |
| --- | --- |
| $\tau$ (ms) | $1.41 \pm 0.13$ |

**Table 1. VADER1's kinetics at 33°C under widefield 1P illumination .**
|  |  |
| --- | --- |
| $\tau_{\text{fast}}$ (ms) | $0.79 \pm 0.10$ |
| $\tau_{\text{slow}}$ (ms) | $20 \pm 8$ |
| % fast | $77 \pm 5$ |

We next quantified VADER1’s ability to report physiologically relevant voltage waveforms. Under resonant 2P scanning, VADER1 detected single action-potential waveforms with ∼40% ΔF/F (Figure 2F, first row; Figure 2G). Because of VADER1’s fast kinetics, the maximal imaging rate of the optical system used for benchmarking (440 Hz) undersampled the response; measured spike amplitudes underestimate the true peak response. Optical signals exhibited a full width at half maximum of ∼6 ms (Figure S2.2A), compared with ∼4 ms for the underlying electrical spike, consistent with modest temporal broadening from indicator kinetics and sampling.

VADER1 also resolved individual spikes within 100-Hz trains and burst waveforms (Figure 2F, second–fourth rows). Consistent with its voltage-response curve, VADER1 responded robustly to spikes and showed small but discernible signals during subthreshold depolarizations (Figure 2F, third row). Under identical imaging conditions, spike-evoked fluorescence changes could not be reliably distinguished for FlicR2, plausibly due to its lower voltage sensitivity (Figure S2.2B,C).

### Trafficking-motif combination drives robust VADER neuronal soma membrane localization

Evaluation of intermediate VADER variants during directed evolution revealed substantial intracellular fluorescence in dissociated rat hippocampal neurons (Figure S2.3A), consistent with prior reports of aggregation or impaired membrane trafficking in GEVIs incorporating anthozoan-derived fluorescent proteins^12,45,46,47^. We thus tested the addition of trafficking motifs, a strategy previously shown to improve membrane targeting in opsins^46^. However, inclusion of commonly used motifs individually—including sequences promoting export from the endoplasmic reticulum^14^ or Golgi apparatus^14,48^—did not substantially reduce intracellular fluorescent puncta (Figure S2.3B–D).

Incorporating the soma-restriction motif from Kv2.1 (“Kv”)^49^ qualitatively increased plasma membrane fluorescence and reduced aggregation, yet a substantial intracellular signal persisted (Figure S2.3E). In contrast, combining the Kv motif with both Golgi- and ER-export tags, arranged in a specific order, yielded efficient and reproducible somatic membrane localization (Figure S2.3H vs G). When expressed in the mouse visual cortex, VADER1 incorporating the composite sequence—called tKv for simplicity—showed qualitatively predominant localization at the plasma membrane of neuronal somata (Figure 2H; Figure S2.4).

### VADER1-tKv reliably reports spikes for one hour in behaving mice

We first evaluated VADER1-tKv in vivo using ultrafast local volume excitation (ULoVE)^50^, a 2P microscopy technique that employs acousto-optic deflectors to record fluorescence signals from single cells with high temporal resolution and sensitivity. This approach ensured a sufficient sampling rate to faithfully capture VADER1-tKv’s rapid spike responses. As a benchmark, we used the green GEVI JEDI-2P-Kv, which produces the largest reported spike amplitudes under ULoVE and has been extensively validated using multiple in vivo 2P optical recording approaches^4,28,25^.

Both indicators were virally expressed in non-overlapping cortical regions within the same mice. Activity was recorded from Layer 2/3 neurons in head-fixed, behaving animals at 7.1 kHz for 10 minutes. Mean imaging depths were comparable (VADER1-tKv: 189 ± 51 µm, mean ± SD; JEDI-2P-Kv: 219 ± 59 µm; Figure S3.1A).

ULoVE resolution enabled detection of GEVI responses to single spikes and spike bursts with high signal-to-noise ratios (Figure 3A). Under these conditions, VADER1-tKv produced responses approximately 2.5-fold larger than those of JEDI-2P-Kv, with amplitudes of 68.9 ± 16.5% ΔF/F versus−28.6 ± 4.1% ΔF/F, respectively (mean ± SD; Figure 3B).

**Figure 3.**
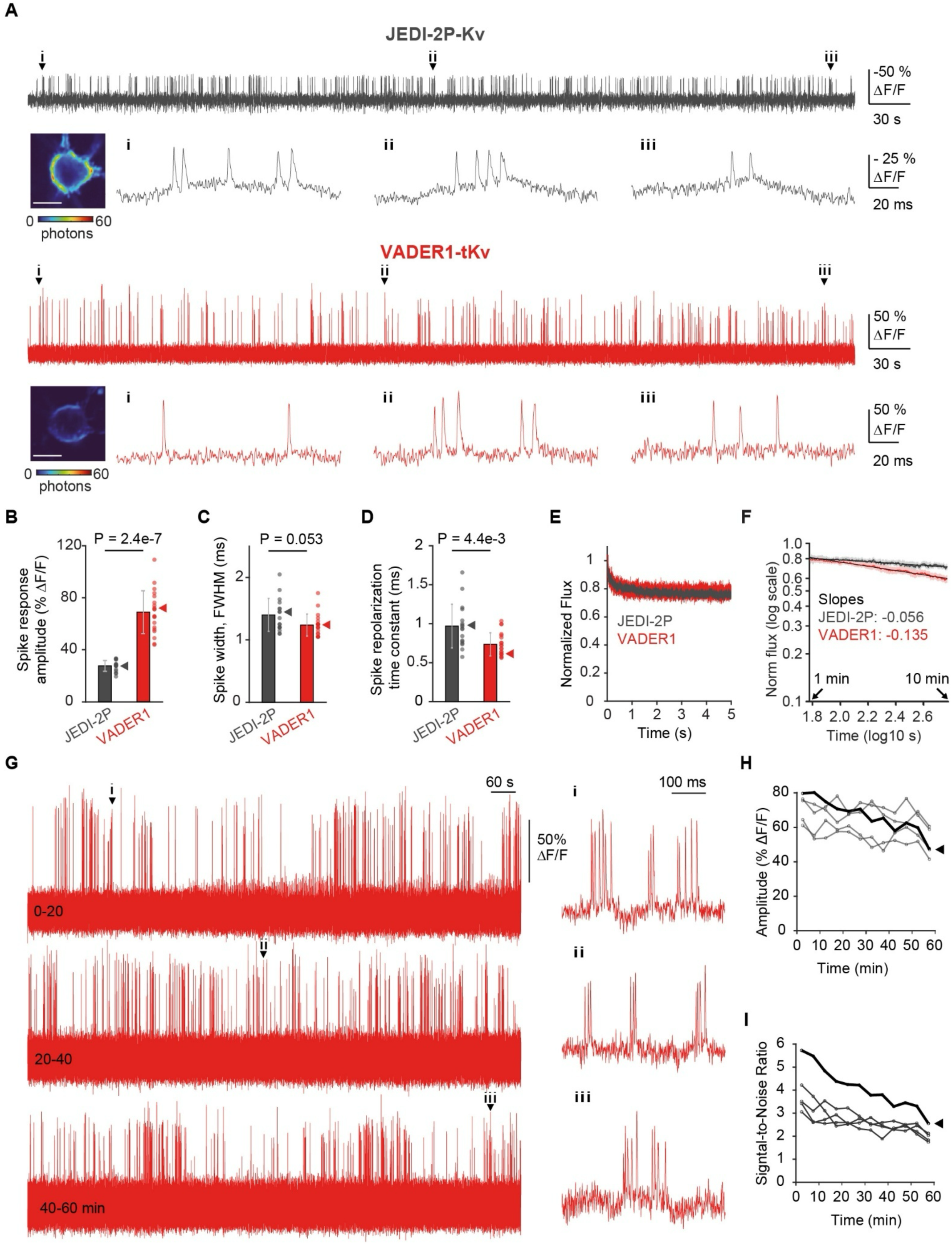
VADER-tKv reports spikes with a larger response amplitude than JEDI-2P-Kv in awake mice. Recordings were acquired at a sampling rate of 7.2 kHz using ULoVE microscopy in layer 2/3 visual cortex neurons transduced with AAVs for Cre-dependent GEVI expression. Excitation wavelength was 1045 nm (VADER1-tKv) and 920 nm (JEDI-2P-Kv). (**A**) Representative 10-min recordings of JEDI-2P-Kv (top, gray) and VADER1-tKv (bottom, red). Power: 22.5 mW measured at the brain surface. The JEDI-2P-Kv trace is inverted to facilitate comparisons. Images of the recorded neurons are shown below the traces (scale bar: 10 μm), followed by randomly selected spike and burst events from the beginning (i), middle (ii), and end (iii) of the recording. A Gaussian filter was applied to the full traces (time width of 0.4 ms) and the zoomed segments (time width of 0.2 ms). Note that scale bars differ between full traces and zoomed segments. (**B**-**D**) Quantitative comparisons. n = 17 (JEDI-2P-Kv) and 20 (VADER1-tKv) cells. Power: 22.5 mW. P values from two-sample Wilcoxon rank sum test. Error bars: SD. (**E**, **F**) Photostability comparison, shown as normalized changes in photon flux during the initial rapid bleaching phase (**E**) and the subsequent slower phase (**F**). Sample sizes are as in (B–D). In (F), dark traces indicate means, shaded regions indicate SEM, and black lines show linear fits. (**G-I**) One-hour recordings from layer 2/3 cortical neurons expressing VADER1-tKv. Excitation power: 18 mW at the brain surface. (**G**) Representative recording. Zoomed segments are shown on the right. The trace was smoothed using a Gaussian filter (0.4-ms width). Additional examples are provided in in Fig. S3.2-3.3. (**H-I**) Change in spike amplitude (**H**) and signal-to-noise ratio (**I**) over the 1-hr recording period. n = 5 cells. The bold trace denotes the cell shown in (G).

Spike responses of VADER1-tKv returned to baseline with a repolarization time constant of 0.74 ± 0.15 ms (mean ± SD), slightly faster than that of JEDI-2P (1.4 ± 0.3 ms, Figure 3D). Despite this difference, the two indicators exhibited statistically indistinguishable optical waveform widths (FWHM, VADER1-tKv: 1.2 ± 0.2 ms, mean ± SD; JEDI-2P-Kv: 1.4 ± 0.3 ms, Figure 3C).

As expected for a positive-going (dim-to-bright) indicator, VADER1-tKv exhibited lower baseline photon flux than the bright-to-dim JEDI-2P-Kv (Figure S3.1B). Nevertheless, signal-to-noise ratio and discriminability were comparable between the two indicators (Figure S3.1C–D; spike SNR: 4.9 ± 1.4 versus 4.6 ± 1.2; D′: 8.8 ± 2.3 versus 9.4 ± 2.7; mean ± SD). Both indicators photobleached biexponentially, with comparable fast bleaching (Figure 3E). Both indicators exhibited biexponential photobleaching with similar fast components (Figure 3E). The subsequent slow phase followed a steeper power-law decay for VADER1-tKv (Figure 3F), yet baseline fluorescence remained at ∼60% of its initial level after 10 min of continuous illumination.

Because the duration of GEVI-based experiments can be limited by photobleaching, we next assessed whether VADER1-tKv could reliably report spikes during a continuous one-hour in vivo session at reduced excitation power (Figure 3G, Figure S3.2). Although spike amplitude and signal-to-noise ratio gradually declined (Figure 3H–I), action potentials remained clearly detectable throughout the full 60 min (Figure 3G-i–iii, Figure S3.3).

Together, these measurements demonstrate that VADER1-tKv combines large spike response amplitudes, fast kinetics, and manageable photobleaching, enabling extended high-speed voltage optical recording in behaving mice with ULoVE microscopy.

### VADER1-tKv enables in vivo spike detection and multiplexed imaging under resonant-scanning microscopy

We next evaluated VADER1-tKv under 2P resonant-scanning microscopy, a widely used method for deep-tissue imaging, albeit with lower temporal resolution than ultrafast methods such as ULoVE. We performed simultaneous voltage imaging and juxtacellular patch-clamp recordings in layer 2/3 of the visual cortex in anesthetized mice across a range of optical conditions (Figure 4A–D; Table S4.1). VADER1-tKv reported individual spikes with a mean response amplitude of 2.3 ± 0.2 σ (mean ± 95% CI; Figure 4C; Figure S4.2); the mean F1 score was 0.49 ± 0.11 (mean ± 95% CI; Figure 4D; Figure S4.2), approaching the value of 0.58 previously reported for JEDI-2P-Kv using the same paradigm^3^.

**Figure 4.**
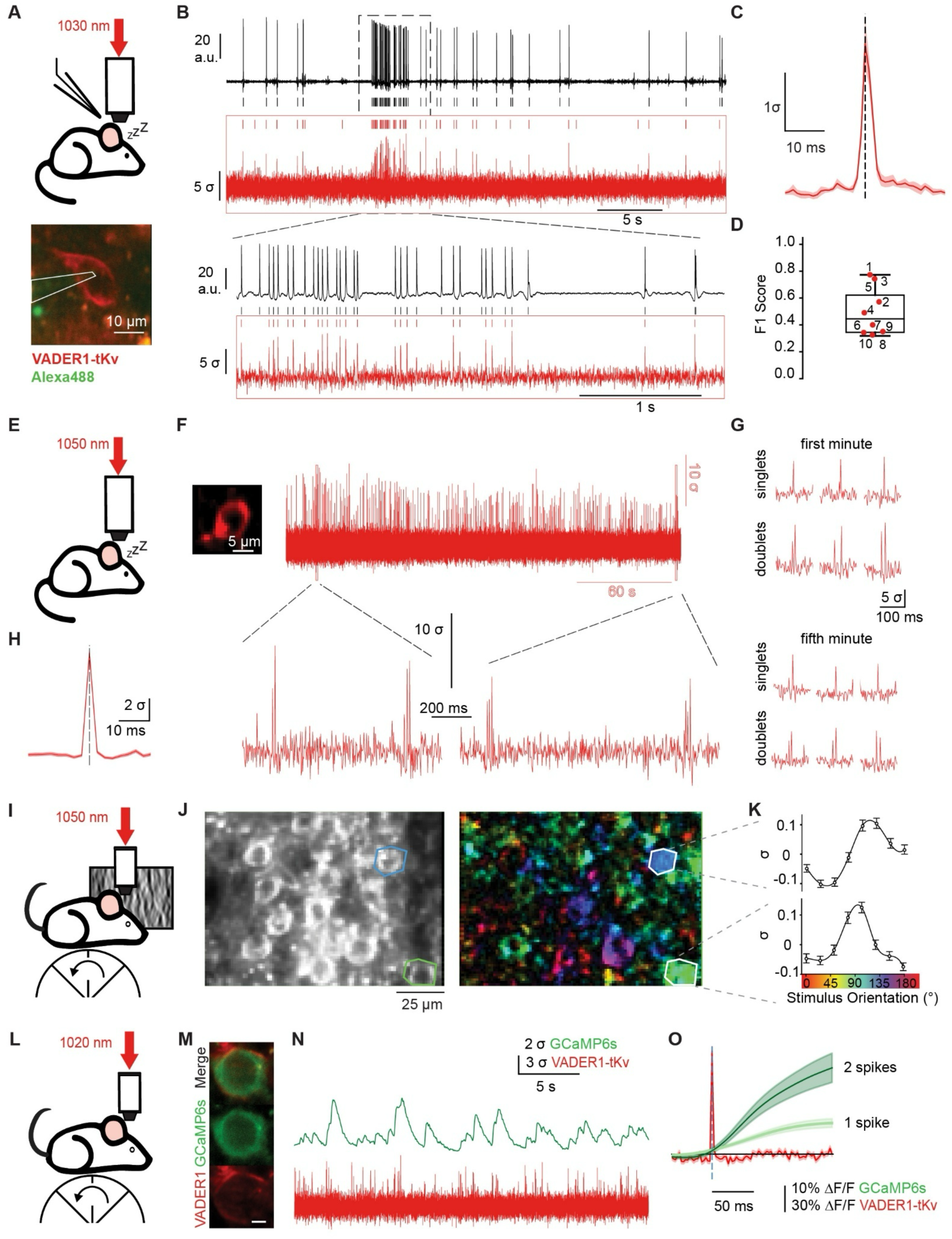
VADER1-tKv enables in vivo action potential imaging and spectral multiplexing under two-photon resonant scanning. All measurements were performed in the primary visual cortex (V1). (**A-D**) In vivo 2P–guided juxtacellular patch-clamp recording combined with VADER1-tKv imaging. (**A**) *Top*: Experimental schematic. Anesthetized animals were used. Imaging depth: 121 ± 32 μm (mean ± SD; range: 69-168 μm) from the brain surface. See Fig. S4.1 for recording details. *Bottom*: high-resolution two-photon image showing VADER1-tKv expression (red) and patch pipette solution (green dye); the pipette outline is indicated. (**B**) Example of simultaneous patch recording (black) and fluorescence imaging (red). Both traces were high-pass filtered at 100 Hz. Frame rate: 720 Hz. Spikes detected using an adaptive threshold are marked by ticks. *Bottom*: Expanded view of the bursting epoch. The trace is from cell 1 (Fig. S4.1). (**C**) Mean VADER1-tKv optical spike waveform obtained by ground-truth spike-triggered averaging. Frame rate: 720 Hz. n = 615 spikes from 6 neurons across 4 mice. (**D**) Spike detection F1 scores, using cell-optimized thresholds. n = 10 neurons from 6 mice. (**E-H**) Imaging spontaneous activity in layer-5 pyramidal neurons. (**E**) Experimental schematic. Anesthetized animals were used. Image acquisition rate: 396 Hz. (**F**) 5-min fluorescence trace of the neuron shown (left). Trace was high-pass filtered at 100-Hz. *Bottom*: Expanded views. Representative trace from n = 4 neurons across 2 mice (other examples provided in Fig. S4.3 and S4.4). (**G**) Example responses to isolated spikes (singlets) and pairs of closely spaced spikes (doublets) during the first (top) and last (bottom) minute of the recording shown in (F). (**H**) Mean VADER1-tKv optical spike waveform obtained by averaging detected spikes. n = 168 spikes. Shaded area: SEM. (**I-K**) Orientation tuning of layer-2/3 neurons in awake behaving mice. (**I**) Experimental schematics. Frame rate: 198 Hz. (**J**) Pixel-wise tuning map (right) for the field-of-view shown on the left. Colors indicate preferred orientation as defined in (K). The tuning map was downsampled for visualization purposes. (**K**) Mean orientation tuning curves for the neurons outlined in (J). n = 200 trials per orientation. Error bars: SEM. (**L-O**) Dual-color, multiplexed imaging of VADER1-tKv and GCaMP6s in awake behaving mice. (**L**) Experimental schematic. Image acquisition rate: 396 Hz. (**M**) High-resolution images showing cellular localization of the indicators. Scale bar: 10 μm. (**N**) Simultaneously acquired fluorescence traces for GCaMP6s (top; 20-Hz low-pass filtered) and VADER1-tKv (bottom, 100-Hz high-pass filtered) for the neuron shown in (M). (**O**) Mean GCaMP6s fluorescence responses aligned to singlet and doublet spikes detected optically with VADER1-tKv. Doublets were defined as two spikes within 50 ms; trace segments were aligned to the first spike. Singlets and doublets were included only if no spikes occurred in the preceding 0.5 s, minimizing contamination from preceding GCaMP6s signals. Red trace: mean VADER1-tKv response to singlets. Traces are filtered as in (N). Shaded areas: SEM.

We next optically recorded neurons continuously for 5 min in layers 2/3 and 5 (Figure 4E–H; Figure S4.3–S4.4). Optical spikes were detected throughout the session, with singlet and doublet events resolved in both early and late epochs, indicating stable high-SNR performance (Figure 4F, 4G). Spike-triggered averaging of isolated singlets yielded a narrow spike waveform, consistent with VADER1’s rapid off-kinetics (Figure 4H).

Functional performance was then assessed in awake animals during presentation of drifting grating stimuli (Figure 4I). Single-pixel analysis is a stringent test, revealing cell-shaped patterns only when responses are robust across independently analyzed pixels. VADER1-Kv showed consistent orientation tuning across pixels within individual neurons, enabling construction of cellular tuning curves (Figure 4J,K).

Finally, to test compatibility for multiplexed imaging, we co-expressed VADER1-tKv with the green calcium indicator GCaMP6s in cortical layer 2/3 (Figure 4L–O). Optical spikes reported by VADER1-tKv coincided with the rising phase of GCaMP6s transients, and pairs of VADER1-tKv-detected spikes elicited larger GCaMP6s responses than singlets (Figure 4O), as expected.

Together, these results demonstrate that VADER1-tKv supports in vivo spike detection and spectrally multiplexed imaging under resonant-scanning 2P microscopy.

## Discussion

VADER1 fills a longstanding gap in the voltage-imaging toolkit by providing a red-shifted, fully genetically encoded voltage indicator that is broadly compatible with a range of 2P microscopy methods and efficiently excited by affordable, high-power ytterbium-doped lasers operating at ∼1030–1064 nm. As a result, VADER1 extends high-speed voltage imaging to a broader range of laboratories and experimental contexts, facilitating wider adoption across the neuroimaging community.

Despite being a first-generation red GEVI, VADER1 exhibits robust in vivo performance. Its signal-to-noise ratio is comparable to that of JEDI-2P, while its response amplitude is considerably larger, even though JEDI-2P represents a more mature, multi-generation indicator. When combined with ultrafast local volume excitation (ULoVE) microscopy, VADER1 supports stable voltage recordings over hour-long sessions, enabling interrogation of neural activity across extended behavioral and physiological time scales.

Beyond single-channel voltage imaging, VADER1 provides a compelling foundation for multiplexed experiments. Its red emission allows straightforward combination with the wide palette of green sensors for neurotransmitter release, neuromodulatory signaling, pH, ions, and metabolism, facilitating multimodal measurements of neural state. Pairing VADER1 with calcium indicators, as we demonstrated, could offer new insights into the nonlinear transformations between voltage and calcium signals^51^, and leveraged to improve calcium-based spike inference. Spectral multiplexing VADER1 with green GEVIs would open the door to population-level comparisons of voltage dynamics across genetically or anatomically defined cell types, and its paring with blue-shifted optogenetic actuators would facilitate 2P all-optical electrophysiology.

Engineering efforts identified several mutations and sequences that provide mechanistic insight and avenues of broader relevance. In particular, the left-shifting mutation located at the C-terminal end of S4 may compensate for overly right-shifted voltage–fluorescence response curves of other (green) GEVIs^52,53^. The tKv combination of trafficking motifs may facilitate efficient plasma membrane expression of other red membrane-based indicators. More generally, VADER1 represents a promising template for continued directed evolution toward indicators that are explicitly optimized for 2P excitation. We do not currently recommend VADER1 for 1P imaging, owing to the complex and suboptimal photophysics of the mApple fluorescent protein under this modality^54,55^.

Priorities for future optimization include additional left-shifting of the voltage response curve to enhance sensitivity to subthreshold voltage fluctuations, including synaptic potentials, which remain largely inaccessible with this first-generation red GEVI. Kinetic tuning is also desired. Faster depolarization kinetics would increase spike-evoked response amplitudes, while moderately slower repolarization kinetics could improve detectability under conventional 2P imaging approaches with limited temporal resolution, such as resonant scanning. Consistent with this interpretation, spike response amplitudes were relatively stable under ULoVE microscopy but showed substantially greater variability under resonant scanning, likely reflecting the mismatch between VADER1’s narrow optical response waveform (∼1.2 ms) and sub-kilohertz sampling rates.

In sum, VADER1 meaningfully advances the ability to perform high-speed, multiplexed 2P voltage imaging in vivo. By expanding spectral access to red, leveraging widely available laser technology, and demonstrating strong first-generation performance, VADER1 lays the groundwork for a new class of 2P-optimized GEVIs. These advances open new opportunities to map functional connectivity and to dissect how voltage dynamics, calcium signaling, and neuromodulatory inputs converge to drive neural computation and behavior.

## Methods

### High-throughput voltage indicator screening

#### Predicted GEVI 3D structures and directed evolution lineage tree

Predicted 3D models of the VADER1 and FlicR2 were generated by AlphaFold3, and rendered using PyMOL (Version 3.0, Schrödinger, LLC). The evolutionary tree of mutations was rendered using Microsoft Office Visio.

#### Reagents

Cloning reagents include PrimeSTAR HS DNA Polymerase (R040A, TaKaRa), Platinum SuperFi II polymerase (12361010, Thermo Fisher Scientific), FastDigest NheI (FD0974, Thermo Fisher Scientific), FastDigest HindIII (FD0504, Thermo Fisher Scientific), FastDigest KpnI (FD0524, Thermo Fisher Scientific), FastDigest Bsp1407I (FD0933, Thermo Fisher Scientific), FastDigest Green Buffer (10X) (B72, Thermo Fisher Scientific), agarose (BP160-500, Fisher BioReagents), Tris-Acetate-EDTA (TAE) 50× Solution (BP1332-1, Fisher BioReagents), Lysogeny broth, Miller (BP1426-500, Fisher BioReagents), Ampicillin (BP1760-25, Fisher BioReagents), In-Fusion HD Cloning Kits (639650, TaKaRa), GeneJET Gel extraction kit (FERK0691, Fisher), 96-well Mini Plus Plasmid Extraction System (Viogene #GF961001), and Mini Plus Plasmid DNA Extraction System (GF2002, Viogene).

Cell reagents include: high-glucose Dulbecco’s Modified Eagle Medium (D1145, Sigma-Aldrich), fetal bovine serum (F2442, Sigma-Aldrich), glutamine (G7513, Sigma-Aldrich), Penicillin/Streptomycin (P4333, Sigma-Aldrich), Geneticin (G418) Sulfate (30-234-CR, Corning), phenol-free Neurobasal medium (12348017, Gibco), B-27 (17504044, Gibco), Glutamax (35050061, Gibco), 30-70 kD poly-D-lysine (P7886, Sigma-Aldrich), ), 300 kD poly-D-lysine hydrobromide (P7405, Sigma-Aldrich), Trypsin-EDTA solution (T3924, Sigma-Aldrich), and Dulbecco’s Phosphate Buffered Saline (Corning, #21-031-CV).

Transfection reagents include JetPRIME (114-15, Polyplus Transfection), FuGENE HD transfection reagent (E2311, Promega), and Opti-MEM (31985-070, Gibco).

Reagents for solutions include: NaCl (S3014, Sigma-Aldrich), sucrose (S0389, Sigma-Aldrich), D-(+)-glucose (G8270, Sigma-Aldrich), HEPES (H3375, Sigma-Aldrich), KCl (P9541, Sigma-Aldrich), MgSO4 (M2643, Sigma-Aldrich), K-gluconate (P1847, Sigma-Aldrich), EGTA (E3889, Sigma-Aldrich), MgCl2 (M9272, Sigma-Aldrich), CaCl2 (223506, Sigma-Aldrich), KOH (P250, ThermoFisher) and NaOH (S5881, Sigma-Aldrich).

### Solutions

- **Growth medium #1**: is comprised of high-glucose Dulbecco’s Modified Eagle Medium supplemented with 10% fetal bovine serum (FBS), 2 mM glutamine, 100 unit/mL penicillin, 100 μg/mL streptomycin, and 750 μg/mL of the antibiotic G418 sulfate (geneticin).
- **Growth medium #2**: is comprised of high-glucose Dulbecco’s Modified Eagle Medium supplemented with 5% FBS, 2 mM glutamine, 100 U/mL Penicillin, and 100 μg/mL Streptomycin.
- **Imaging solution #1**: is comprised of 110 mM NaCl, 26 mM sucrose, 23 mM glucose, 20 mM HEPES, 5 mM KCl, 2.5 mM CaCl_2_, 1.3 mM MgSO_4_, adjusted to pH 7.4 with NaOH, and adjusted to 300 mOsm/kg with H_2_O.
- **Imaging solution #2**: is comprised of 106.5 mM NaCl, 26 mM sucrose, 23 mM glucose, 20 mM HEPES, 8.5 mM KCl, 2.5 mM CaCl_2_, 1.3 mM MgSO_4_, adjusted to pH 7.4 with NaOH, and adjusted to 300 mOsm/kg with H_2_O.
- **Internal solution:** is comprised of 115 mM K-gluconate, 10 mM HEPES, 10 mM EGTA, 10 mM glucose, 8 mM KCl, 5 mM MgCl_2_, 1 mM CaCl_2_, adjusted to pH 7.4 with KOH, and adjusted to 290 mOsm/kg with H_2_O.

### Library construction

GEVI libraries were generated by site-directed saturation mutagenesis targeting single residues. To obtain a more uniform distribution of residues for saturation mutagenesis, we combined primers containing the NNT, VAA, ATG, or TGG codons (N = any base; V = A, G, or C) at a molar ratio of 16:3:1:1. The 20-μL PCR reaction mix contained 1 μL of a 10-μM forward primer mix, 1 μL of a 10-μM reverse primer, 5 ng of template plasmid, and 10 μL of a 2× PCR master premix (PrimeSTAR HS DNA polymerase, Takara). DNA was amplified using the following protocol: an initial denaturation step at 98°C for 30 s; 30 amplification cycles consisting of 98°C for 10 s, 57°C for 10 s, and 72°C for 1 minute/kb of fragment length; and a final extension step at 72°C for 5 minutes. The pcDNA3.1/Puro-CAG^1^ backbone was linearized using the restriction enzymes NheI and HindIII. PCR products and linearized backbones were purified using gel electrophoresis and the GeneJET Gel Extraction Kit. PCR products, including a C-terminus P2A-linked EBFP2-CAAX reference protein^2^, were assembled with the vector backbone using the In-Fusion assembly system according to the manufacturer’s instructions. The In-Fusion reaction mix was transformed into commercial chemically competent bacteria (XL10-Gold, Agilent) with a transformation efficiency exceeding 5×109 FU per μg of DNA. Liquid cultures were inoculated with manually picked colonies, and purified plasmids were prepared using a 96-well plasmid purification kit following the manufacturer’s instructions.

### Plasmid construction for in vitro characterization

Plasmids used for in vitro characterization were assembled using the In-Fusion cloning technique into a pcDNA3.1/Puro-CAG vector with or without a C-terminus P2a-EBFP2-CAAX reference protein. Liquid cultures were inoculated with manually picked colonies, and purified plasmids were prepared using a purification kit following the manufacturer’s instructions. The plasmid sequence was confirmed using whole-plasmid nanopore sequencing (Plasmidsaurus Inc.) or Sanger sequencing (Eurofins Genomics LLC). The FlicR2 sequence was amplified from the pCMV-FlicR2 plasmid, a gift from Dr. Madhuvanthi Kannan. The mApple sequence was amplified from the pCMV-mApple plasmid, a gift from Dr. Koen Venken’s lab.

### Cell culture and transfection in 96-well plates

To screen GEVIs for responses within the physiological range, we used HEK293 cells stably expressing the human Kir2.1 channel^3^, which maintains a resting membrane potential of approximately −83 mV using imaging solution #1. This membrane potential was chosen because it approximates the lower bound of the physiological range for mammalian neurons, enabling screening for improved responses across the hyperpolarized state and into the depolarized range.

HEK293-Kir2.1 cells were cultured at 37 °C with 5% CO2 in growth medium #1. G418 was included to maintain the expression of the Kir2.1 transgene, which was chromosomally integrated with a G418 resistance gene. For screening GEVIs, glass-bottom 96-well plates (P96-1.5H-N, Cellvis) were first coated with poly-D-lysine (30-70 kDa) to promote cell adherence to the glass. The coating was performed for 1 hour at room temperature or 37 °C, and the plates were washed twice with DPBS. HEK293-Kir2.1 cells were then plated to 60-80% confluency in 100 μL of growth medium #2.

We randomly selected 88 variants per saturation mutagenesis library. According to statistical modeling^4^, our library generation and sampling strategy produced a ∼99% theoretical probability that any given screened library included the best residue. Each well was transfected with jetPRIME (Polyplus) according to the manufacturer’s protocol, using the following specifications: a mixture of 130 ng DNA, 0.4 μL jetPRIME transfection reagent, and 10 μL jetPRIME buffer was combined with 50 μL growth medium #2, then added to the well. Independent transfections were defined as those in which DNA was added to each well separately. After 6-18 hours, 100 μL growth medium #2 from each well was replaced with fresh growth medium #2 to minimize potential cytotoxicity from the transfection reagents. Two days post-transfection, the cells were washed twice with 200 μL of imaging solution #1 at room temperature. Wells were filled with 100 μL of the imaging solution #1 and screened with our high-throughput screening platform (see below). The imaging solution #1 was adjusted with H_2_O to achieve a final osmolarity of 290-310 mOsm/kg.

### RFP/OFP membrane localization screening

The following RFPs and OFPs were inserted into a voltage-sensing domain and screened for membrane localization: LSSmCrimson^5^, TagRFP-T^6^, mRuby4^7^, mCherry^8^, mTagRFP^9^, mRuby3^10^, mScarlet-I^11^, mBanana^8^, mStrawberry^8^, mApple^12^, and mOrange2^12^. FPs were inserted between the 3^rd^ and 4^th^ transmembrane helices using the cloning strategy described in the section “Plasmid construction for in vitro characterization”. HEK293-Kir2.1cells were cultured and transfected in 96-well plates with each construct as described in the section “Cell culture and transfection in 96-well plates”. Single red-channel images were taken for each construct using the same microscope and optical setup as described in the section “high-throughput screening under one-photon”, except that the FOV size was 1024 × 1024 or 2048 × 2048 pixels and the exposure time was 20 ms.

Raw Nikon .nd2 images were converted to single-channel TIFF files using Bio-Formats and analyzed in MATLAB (R2023a, MathWorks) with custom scripts. Images were background-corrected before analysis. Cell segmentation masks were generated using Cellpose and imported as labeled masks. Individual ROIs were filtered to exclude objects with areas <500 or >1500 pixels or that contained saturated pixels. For each ROI, a narrow band corresponding to the membrane was defined by dilating the ROI perimeter and intersecting it with the ROI mask. The cytoplasmic region was defined as the remaining interior ROI pixels excluding the membrane band. Mean fluorescence intensities were computed for the membrane (*F_mem_*) and cytoplasmic (𝐹*_cyt_*) compartments, and a membrane localization metric was calculated per cell as the membrane-to-cytoplasm ratio (*F_mem_*/𝐹*_cyt_*).

For visualization and quality control, a one-dimensional fluorescence profile across each cell was extracted by generating a line intersecting the cell boundary using a Hough transform; the line whose midpoint was closest to the ROI centroid was selected, and a perpendicular line through this midpoint was used to sample fluorescence intensity across the cell. Profiles were excluded if the sampled line length was outside a predefined range corresponding to approximately one cell diameter, with thresholds scaled according to image resolution (1024 × 1024 vs. 2048 × 2048), or if the normalized intensity profile contained NaN values. These profiles were used solely for quality control and were not involved in membrane localization quantification.

Membrane-to-cytoplasm ratios were averaged per construct within each imaging dataset and normalized to the mean value of the mCherry-CAAX control measured in the same dataset. Normalized per-cell values were then pooled across datasets for each construct and reported as relative membrane localization.

### High-throughput GEVI screening platform

To evaluate the performance of GEVIs at high throughput, we built an automated multimodal 96-well screening platform based on an inverted microscope (Ti, Nikon Instruments). The excitation light was generated from a laser diode light engine (LDI, 89 North) and directed to the microscope’s epifluorescence illuminator via a liquid light guide. Violet (405 nm) and green (555 nm) light were delivered to the sample plane through a 20× 0.75 NA objective (CFI Plan Apochromat Lambda, Nikon Instruments) to excite the EBFP2 and mApple-based GEVIs, respectively. The emission light from the sample plane was split from the excitation light through a quad-band dichroic mirror (ZT 405/470/555/640rpc-UF2, Chroma) and filtered by a quad-band emission filter (ZET 405/470/555/640m-OD8, Chroma) before being collected by a scientific complementary metal–oxide–semiconductor (sCMOS) camera (either ORCA Flash 4.0 V2, C11440-22CU, Hamamatsu, or a Kinetix, Teledyne Vision Solutions). A motorized extended-travel stage capable of holding 96-well plates (TI-S-ER, Nikon) was used to control the field-of-view position.

To support system automation, data-acquisition and output boards (PCI-6723, National Instruments) were connected to the microscope computer via a PXI Chassis (PXI-1073, National Instruments). The computer was equipped with an Intel Xeon W5-3435XE5 processor (16 cores), 256 GB of DDR4 RAM, and eight 2 TB SSDs in RAID 0 (SSD7540 PCIe 4.0 × 16 8-Port M.2 NVMe RAID Controller, Highpoint Technologies) to facilitate high-speed imaging. JOBs scripts in NIS-Elements HC (version 5.42.06, Nikon Instruments) were used to control the microscope system (e.g., stage position), manage optical configurations (e.g., excitations), initiate image acquisition, and trigger the stimulator.

A digital, isolated, high-power stimulator (4100, A-M System) was used to provide electric field stimulation. Electric pulses were passed through a pair of electrodes made from 0.5 mm-wide platinum wires (99.95% pure, AA10286BU, Fisher Scientific). The two L-shaped electrodes had a horizontal length of 2 mm and were spaced 3 mm apart (Figure 1B) and were secured to a 3D-printed polylactic acid holder. The holder was fixed to a motorized linear translation stage (LSM050B-E03T4A, Zaber), controlled by Zaber Launcher (Zaber) to move the electrodes in and out of individual wells. Two manual linear translation stages (411-05S, Newport) were used to fine-tune the lateral position of the electrodes. During stimulation, the electrodes were submerged under the imaging solution, about 500 μm above the cells.

### High-throughput GEVI screening under widefield one-photon illumination

For variants VADER0.1 through VADER0.4, four non-overlapping fields of view (FOVs) of 2048 × 1024 (512 × 256 after applying 4 × 4 binning) pixels were imaged per well with a sCMOS camera (ORCA Flash 4.0 V2, C11440-22CU, Hamamatsu). The electric field stimulation (EFS) electrode pair was moved into the well after the motorized stage centered at the well and before the stage moved to the first selected FOV. The reference channel for EBFP2 brightness was captured first under 405 nm illumination for 1 frame, with an irradiance of ∼11 mW/mm^2^ at the sample plane and an exposure time of 10 ms. Then, the target channel designated to capture responses from the red GEVIs was imaged at ∼200 fps under 555 nm illumination for 900 frames (∼4.5 s), with an irradiance of ∼ 412 mW/mm^2^ at the sample plane. Immediately before the camera started, a transistor-transistor logic (TTL) signal was sent to trigger the stimulator, which sent out a TTL signal to trigger the shutter of the light source and turn the light on. This triggering allowed the camera to capture the onset of the excitation light, which eventually served as a reference point for locating EFS events and temporally aligning results across different FOVs. One second after the light was turned on, field stimulation pulses were applied. The field stimulation train consisted of three monophasic 60-V, 1-ms pulses, followed by three 20-V, 20-ms pulses, with a 500-ms inter-pulse interval. The train finished with a 200 Hz burst of 20 30-V, 1.5-ms pulses. After the EFS on the 4 selected FOVs, the motorized stage was re-centered in the well, and the motorized linear translation stage raised the electrode above the well.

For screening VADER0.5 through VADER1, the same procedure was followed, with the following changes: 1200 × 520-pixel FOVs (no binning) were imaged at ∼510 fps for 5000 frames (∼9 seconds) with a Kinetix sCMOS camera (Kintex Teledyne Vision Solutions) mounted on a 0.45× demagnifying tube (Nikon). The field-stimulation protocol began with five monophasic square pulses, each with a 1-ms width, 60-V amplitude, and an inter-pulse duration of 300 ms. These pulses were followed by a series of seven 2.5-ms width pulses spaced 300 ms apart with increasing voltage amplitudes (5, 10, 15, 20, 25, 30, and 35 V), then a 100-Hz train (5 monophasic square pulses with 2.5-ms width, 30-V amplitude, and an inter-pulse duration of 10 ms), and finally another five 1-ms, 60-V amplitude pulses.

### High-throughput screening data analysis

Image analyses were performed using in-house developed Python software (Python 3.12.10). Whole-cell segmentation was performed using Cellpose 3(cellpose reference). Background fluorescence for both GEVI and reference channels was estimated as the median intensity of the inverse cell mask, excluding all cell-containing regions. Cells were excluded from analysis if their mean fluorescence intensity did not exceed two standard deviations above the mean background signal or if more than 5% of pixels within the cell were saturated at any point during acquisition. To restrict measurements to the plasma membrane, a Laplacian filter was applied, followed by an inward-directed Gaussian blur corresponding to a width of 1 μm.

Photostability was quantified using normalized fluorescence time courses excluding stimulus-associated frames. For each cell, fluorescence intensity was normalized to the maximum observed signal before fitting. Normalized traces were fit to single-, double-, or triple-exponential decay models to account for heterogeneous bleaching kinetics across cells, and model selection was performed using Akaike and Bayesian information criteria, selecting the model with the highest shared likelihood score^13^. Photostability was defined as the area under the curve (AUC) of the selected fit, normalized to the maximum possible AUC given the total acquisition time, with a value of 1 indicating no detectable photobleaching. Brightness was defined as the ratio of mean GEVI to mean reference protein fluorescence intensity following photobleaching correction. Brightness dynamics were calculated as the difference between the GEVI-to-reference ratio measured in the bright and dim states of the GEVI of interest. Response magnitude was defined as the peak fluorescence change evoked by a given stimulation pulse. All metrics were calculated at the single-cell level and averaged within each well, with wells treated as independent biological replicates for data representation.

Compound metrics that combine multiple performance metrics were used to simplify indicator ranking^14^. We previously developed the detectability index metric (𝐷_’_ = 𝑅√𝐵, arb. unit) — where 𝑅 is the peak response amplitude and 𝐵 is the relative brightness. We also sought to consider the impact of photobleaching on voltage recording and avoid variants that are bright but bleach rapidly. Thus, we used the detectability budget (𝐷_B_ = 𝐷*_I_*√𝑃, arb. unit)—which combines the peak response amplitude 𝑅, the relative brightness 𝐵, and the photostability 𝑃. None of the above metrics include physical units, as they are relative parameters intended solely for comparative ranking of indicators during the screening process.

To evaluate the reliability of well-level analysis, we applied a rank-based normalization approach to ensure comparability across metrics within a given construct. For each metric, values were converted to percentile ranks. An aggregate performance score was computed as the mean percentile, penalized by within-row variability (standard deviation). A consistency weight of 0.5; data points falling within the extreme tails of the distribution (≤1st percentile or ≥99th percentile) were flagged as potential under- or over-performers. To ensure that exclusions reflected true multivariate abnormalities rather than metric-specific noise, flagged candidates were subsequently validated using both Grubbs’ test for univariate outlier detection and Mahalanobis distance to assess deviation in the joint feature space. Only data points that met both rank-based extremity criteria and statistical outlier significance under these tests were excluded from downstream analysis.

### Membrane localization tag screening in dissociated cortical neurons

E18 mouse cortical neurons were purchased from Transnetyx tissue by BrainBits (SKU SDEDHP, Transnetyx, Inc.) and came in Hibernate® EB complete Media (BrainBits HEB-100 mL) as single-cell suspensions. Before seeding the hippocampal neurons, a 24-well glass-bottom plate (P24-1.5H-N, Cellvis) was coated with Neuron Coating Solution (027-05, Sigma-Aldrich) at 37 °C overnight and then washed three times with DPBS. Neurons were seeded at 80,000–100,000 cells/mL in 500 µL glutamate-supplemented NbActiv1 medium (NB1 + GLU, Transnetyx Inc.) per well of a 24-well plate, following the manufacturer’s protocol. The day of plating was considered day in vitro (DIV) 0. Culture medium was replaced with pre-equilibrated NbActiv4 medium (SKU NB4, Transnetyx Inc.) on DIV 1, and half of the media was replaced every 3-4 days thereafter.

Transfection was conducted on DIV 6 using calcium phosphate, as previously described^15^. In short, calcium phosphate-DNA precipitates were prepared by first combining plasmid DNA (a mix of 150 ng pAAV-hSyn-VADER0.2-[*membrane localization tag*], 50 ng pCaggs-EBFP2-CAAX, and 600 ng pNCST bacterial expression vector acting as buffer/filler DNA (pNCST backbone is based on Addgene plasmid #129508) with 18 µL sterile water, followed by the addition of 1.25 µL 2.5 M CaCl_2_. The mixture was gently pipetted and incubated at room temperature for 1 hour. Conditioned medium was collected and saved. Neurons were briefly washed with warm NbActiv4 medium, and calcium phosphate-DNA precipitate was added dropwise to each well. Plates were gently shaken and returned to the incubator for 2 hours. Immediately before the end of incubation, the 12.5 mL NbActiv4 medium aliquot was acidified with 12 N HCl until the medium turned yellow. The transfection medium was replaced with acidic NbActiv4 medium for 5–8 minutes at room temperature until the precipitate dissolved. Acidic NbActiv4 medium was then replaced with fresh 37 °C NbActiv4 medium, followed by restoration of the original conditioned medium.

A week after transfection (DIV 13–15), the attached neurons were washed twice with imaging solution #1, then 500 µL of imaging solution #1 was added to each well. Single images of the red and blue channels were acquired using the same microscope and optical setup as described in the section “High-throughput screening under one-photon”, except that the FOV was 1024 × 1024 pixels.

### In vitro characterization of VADER

#### One-photon excitation and emission spectra characterization

To characterize the one-photon excitation and emission spectra of GEVIs, we used the same vector used for in vitro GEVI characterization with whole-cell voltage clamp, i.e., pcDNA3.1/Puro-CAG-*GEVI*, and a control vector, pcDNA3.1/Puro-CAG-mApple. Transfections were performed in 10-cm tissue culture dishes (353003, Corning), and HEK293A cells (Thermo Fisher Scientific) were seeded one day before forward transfection at 60–65% confluency, equivalent to 720,000–780,000 cells per well or 5,280,000–5,720,000 cells per dish in growth medium #2. Transfections were performed 36–48 h before imaging using JetPRIME. The transfection medium was replaced 4 h after transfection with fresh growth medium #2 to minimize potential cytotoxicity from the transfection reagent.

On the day of the experiment, cells from a 10-cm dish transfected with the same construct were detached with trypsin, washed twice, diluted into imaging solution #1, and evenly split into three wells of a glass-bottomed 96-well plate (P96-1.5H-N, Cellvis). Pooling the cells into a dense preparation was essential to produce a strong signal that the plate reader could robustly detect. Untransfected cells were prepared with the same method and deposited in a separate well to determine the background autofluorescence levels. A handheld automated cell counter (Scepter 3.0, Millipore) was used to plate a similar number of cells in each well.

Spectra were determined using a plate reader (Cytation 5, BioTek) to quantify fluorescence from wells of the 96-well plates prepared above. Emission spectra were acquired by exciting at 525/10 nm and measuring emitted photons from 550 to 700 nm in 1-nm increments, with a 10-nm bandwidth. Excitation spectra were acquired by exciting from 350 to 628 nm in 1-nm increments, with a 10-nm bandwidth, and measuring emission at 650/10 nm. Individual spectral scans were corrected for autofluorescence by subtracting the values from untransfected cells at each wavelength and then normalized to their respective peaks. The final emission spectra were obtained by averaging the normalized spectra from each scan. The resulting spectra were smoothed using the Savitzky-Golay method^16^ employing a second-order polynomial with 4 neighbors on each side, and then normalized to the peak intensity. Peak emission wavelengths were determined by averaging the peak excitation/emission wavelengths across all scans.

#### Two-photon excitation spectra

CHO cells were acquired from Sigma (Sigma, 85050302) and cultured in T25 flasks (Falcon, 353107) in a medium consisting of DMEM-F12 + Glutamax (Fisher, Gibco™ 10565018), supplemented with 10% FBS (Fisher, Gibco™ 10500064) and 1% penicillin/streptomycin (5000 U·mL−1). Cells were passaged every 2–3 days, until P20, to avoid genetic drift. Before each experiment, cells were seeded on glass coverslips (Fisher, 10252961) in a 35 mm petri dish (25,000 cells/mL). After 24 hours, cells were transiently transfected with either pcDNA3.1/Puro-CAG-VADER1, pcDNA3.1/Puro-CAG-mApple, or pcDNA3.1/Puro-CAG-FlicR2 plasmid using the Jet prime kit (Ozyme, POL101000015). Experiments were performed 48−72 hours post-transfection on live CHO cells at room temperature in a HEPES-buffered saline solution (pH adjusted to 7.4).

Two-photon spectra were measured using a scanning microscope (Independent NeuroScience Services, INSS Ltd, UK) coupled to two tunable lasers: (1) Ti:sapphire laser (80 MHz, 100 fs, Chameleon Ultra, Coherent) used at 800-1080 nm and (2) an optical parametric amplifier tunable source (I-OPA-TW-HP, 0.5 MHz, 180 fs, Light Conversion) used at 1070-1300 nm pumped by a low-rep femtosecond laser (Carbide, 0.5 MHz, 210 fs, Light Conversion). Fluorescence images of expressing cells were collected by scanning across a field of view of ∼300 × 300 µm with a 16× 0.8NA objective (Nikon) and detected by two PMTs (630/75 nm in red and 510/80 nm in green). Fluorescence intensities were measured at constant power across the spectrum. The power of the two lasers was adjusted for the difference in pulse width and repetition rate between the lasers so that they induced the same two-photon excitation effect: P1 = 9.4 ± 0.05 mW for Ti:sapphire (from 800 to 1080 nm) and P2 = 1 ± 0.005 mW for I-OPA-TW-HP (1070-1300 nm). To mitigate chromatic focal drift, z-stacks of 25 µm were acquired. Bleaching was assessed by repeating recordings at 1070 nm at regular intervals, with no significant difference in average intensity.

Fluorescence was measured as the average pixel intensity of manually-segmented expressing cells. We used maximum intensity projections of the acquired stacks and subtracted the local background, normalizing each cell to its own peak intensity. Cells were considered for analysis based on their morphology (roundness) and the absence of green autofluorescence. Outlier spectra with individual point values exceeding twice the standard deviation at any given wavelength were excluded.

#### Brightness under two-photon illumination

To determine the brightness under 2P illumination, HEK293-Kir2.1 cells were transfected with pcDNA3.1/Puro-CAG-*GEVI*-P2a-EBFP2 plasmids and prepared for imaging, as described in “Cell culture and transfection in 96-well plates”. However, imaging solution #2 was used instead of imaging solution #1 to obtain a resting membrane potential of ∼-70 mV, approximating the resting voltage of mammalian cortical pyramidal neurons. Images of the cells were acquired using an inverted microscope with multiphoton capability (A1R-MP, Nikon Instruments) with a 20× NA 0.75 objective (CFI Plan Apochromat Lambda, Nikon). Red GEVIs were excited with 189 mW of 1050 nm light generated by a fiber laser (Ultra FF 1050, Toptica) and blue reference EBFP2 was excited with 93 mW of 750 nm light generated by a Ti:sapphire femtosecond laser (Chameleon Ultra II, Coherent). Laser power was set using an acousto-optic modulator and delivered to the sample plane through a 20× 0.75-NA objective (CFI Plan Apochromat Lambda, Nikon Instruments). Four 1024 × 1024-pixel FOVs were captured for each well using two galvanometer optical scanners with a dwell time of 12.1 µs. Red (GEVI) and blue (EBFP2) images were captured separately. A 490 LP dichroic mirror was used to separate the red and blue channels, followed by either a 605/70 nm (GEVI) or 440/80 nm (EBFP2) emission filter. Emitted photons were detected by gallium arsenide phosphide (GaAsP) photomultiplier tubes (PMTs). A motorized travel stage (H139E1, Prior or SCANplus IM 130 × 85 - 2mm, Märzhäuser Wetzlar GmbH & Co. KG) was used to control the field-of-view position and to hold 96-well plates during imaging.

To support system automation, data-acquisition and output boards (PCI-6229 and PCI-6723, National Instruments) were connected to the microscope computer via a PXI Chassis (PXI-1033, National Instruments). The computer was equipped with an Intel Xeon Gold 6334 processor with 8 cores and 3.60 GHz, 256 GB of DDR4 RAM, and two 12 TB 7200 RPM hard drives configured in RAID 0 to facilitate high-speed imaging. JOBs scripts in NIS-Elements HC (version 5.42.07, Nikon Instruments) were used to control the microscope system (e.g., stage position), manage the optical configurations (e.g., excitations), initiate image acquisition, and trigger the stimulator.

Images were background-subtracted, and the relative brightness was then calculated from the foreground mask as the average pixel intensity in the GEVI channel divided by the average pixel intensity in the EBFP2 channel.

#### Whole-cell voltage clamp setup

HEK293A cells were plated on 30-70 kDa poly-D-lysine-coated circular cover glass (12 mm, #0, 633009, Carolina) at 30% confluence in growth medium #2, two days before imaging. Transfection was performed on the same day as plating, using 100 ng of DNA and 0.3 μL of FuGENE HD (Promega Corporation) transfection reagent per well of a 24-well plate (Costar, #3524), following the manufacturer’s instructions. Unless indicated otherwise, pcDNA3.1/Puro-CAG plasmids expressing the GEVI with no reference protein were used. The cells were cultured at 37 °C with 5% CO_2_ before and after transfection. The next day, the transfection medium was replaced with fresh growth medium #2 to minimize potential cytotoxicity from the transfection reagent.

Glass micropipettes (1B150-F-4, World Precision Instruments) were pulled using a pipette puller (P1000, Sutter) and had tip resistances of 2-6 MΩ. Micropipettes were loaded with internal solution. The internal solution was adjusted with H_2_O to be 10 mOsm/kg lower than the imaging solution #1, which typically resulted in an internal solution between 280 and 300 mOsm/kg. The micropipette was mounted on a patch-clamp headstage (CV-7B, Molecular Devices) and positioned using a micromanipulator (SMX series, Sensapex or a GmbH Mini23, Luigs & Neumann). The coverslip seeded with the transfected cells was placed in a custom glass-bottom chamber designed for the Chamlide EC (Live Cell Instrument), with the bottom of the chamber made from a 24 × 24 mm #1 glass coverslip (Thermo Scientific). Cells were continuously perfused with an imaging solution #1 at approximately 4 mL/minute using a peristaltic pump (505DU, Watson Marlow). Whole-cell voltage-clamp was performed using a MultiClamp 700B amplifier (Molecular Devices). Patch-clamp data were recorded using an Axon Digidata 1550B1 Low Noise system with HumSilencer (Molecular Devices). Command voltage waveforms were compensated for the liquid junction potential of -11 mV. Recordings were considered satisfactory and included in the final analysis only if, both before and after the recording, the patched cell had an access resistance (𝑅*a*) < 12 MΩ and a membrane resistance (𝑅*m*) > 10 × 𝑅*a*.

### Simultaneous electrophysiology and one-photon illumination recordings

#### Evaluating indicators’ voltage response curves under one-photon illumination

To evaluate the sensors’ response to steady-state voltages, we simultaneously illuminated and voltage-clamped cells expressing pcDNA3.1/Puro-CAG-*GEVI* in HEK293A cells. The same patching hardware setup described in “Whole-cell voltage clamp setup” was used, and the same imaging hardware described in the “High-throughput GEVI screening platform” section was used.

The voltage-clamped cell was centered within a FOV of 2048 × 200 (512 × 50 after 4 × 4 binning) pixels. Cells were illuminated by 555 nm, 280 mW/mm^2^ light from an LDI light engine (89 North) and recorded at room temperature. The emission light from the cell was filtered with a multi-band emission filter (ZET405/470/555/640m-OD8, Chroma) and collected by the ORCA Flash 4.0 V2 sCMOS camera at ∼987 Hz with an averaged exposure time of 1.01 ms. We applied 1-s step-voltage stimulations of −100, −80, −60, −40, −20, 0, 20, 30, and 50 mV to the voltage-clamped cells, with a 1.5-s gap at –70 mV between each pair of stimulations, for VADER0.2 and VADER0.3. For VADER0.4, VADER0.5 and VADER1, an extended voltage stimulation set was used (−100, −80, −60, −40, −20, 0, 20, 30, 50, 70, 90, and 120 mV)

Time series recordings from the camera were processed using a method similar to that for high-throughput electric field stimulation recordings (see “High-throughput screening data analysis”).

#### Kinetics

Kinetics were evaluated under one-photon illumination. The same imaging hardware setup described in the “High-throughput GEVI screening under widefield one-photon illumination” section was used, with the following modifications. Cells were illuminated with 555-nm light (SpectraIII, Lumencor) and conditioned using the 440-570-nm band of a multi-band dichroic mirror (77015970, Semrock). The irradiance at the sample plane was ∼0.6-1.3 mW/mm^2^. Green-emitted photons were reflected towards a multialkali photomultiplier tube (PMT, PMM02, Thorlabs) installed on one of the side ports of the microscope using the 578-611-nm band of the multi-pass dichroic (77015970, Semrock) and then filtered at 580-610 nm using a multi-pass filter (77015970, Semrock).

Kinetics were evaluated using three 1-s 100-mV depolarization pulses from −70 to 30 mV. Between each pulse, cells were held at –70 mV for 1.4 s. Recordings were performed at 32-33°C using a feedback-controlled inline heater system (inline heater SH-27B, controller TC-324C, cable with thermistor TA-29, Warner instruments) to maintain the temperature in the perfusion chamber. A diaphragm was used to reduce the diameter of the excitation spot so that during imaging, only one cell at the center of the FOV was illuminated. To maximize photon collection, an oil-immersion 40× NA 1.3 objective (CFI Plan Fluor DIC H/N2, Nikon Instruments) was used. A custom MATLAB (R2023a, MathWorks) routine was used to control the PMT bias voltage and simultaneously record the command voltage and PMT output via a National Instruments data acquisition system. The PMT gain was modulated through an analog output channel, while voltage signals were acquired through analog input channels in a trigger-synchronized manner. Data were sampled at 80 kHz and saved for offline analysis.

The PMT output voltage was analyzed using a custom MATLAB (R2023a, MathWorks) routine to obtain the fluorescence signal for each cell. The raw data was first downsampled to 20 kHz. Then, the photobleaching correction was performed by fitting the baseline (when the cell was held at −70 mV) with a three-term exponential and removing the trend from the entire signal using division. The corrected signal was cropped from 0.1 s before the estimated depolarization or repolarization onset to 1 s after the estimated depolarization or repolarization onset. The exact onset timing was fitted together with other coefficients with either single-exponential (**F(t)** = *c* + (*k* × exp((**t**—*t_0_*) × *λ*)) × (**t** > *t_0_*) + *k* × (**t** <= *t_0_*)) or dual-exponential ((**F(t)** = *c* + (*k* × exp((**t**—*t_0_*) × *λ*) + *k_2_* × exp((**t**—*t_0_*) × *λ_2_*)) × (**t** > *t_0_*) + (*k* + *k_2_*) × (**t** <= *t_0_*)) model where the **t** is the independent variable, **F** is the dependent variable, and the rest of the parameters are coefficients to be fitted. Among these coefficients, *c* describes the mean plateau fluorescence, *k* and *k_2_* describe the relative ratio of each exponential component, *λ* and *λ_2_* describe the opposite reciprocal of time constants, and *t_0_* is an offset indicating the exact event onset timing.

### Simultaneous electrophysiology and two-photon illumination recordings

To achieve sufficient expression, stable HEK293a cell lines expressing VADER1 and FlicR2 were generated, followed by transfection with the pcDNA3.1/Puro-CAG-GEVI plasmid. The same imaging hardware setup was used as described in the “Brightness under two-photon illumination” section. An oil-immersion 40× NA 1.3 objective (CFI Plan Fluor DIC H/N2, Nikon Instruments) was used to focus 1050 nm light set to ∼45 mW onto the sample plane. Videos were taken with a resolution of 512 × 32 pixels and a frame rate of 440 Hz under resonant scanning microscopy. Recordings where noise was indistinguishable from the signal due to the low brightness of the cell were removed from the final analysis. Electrophysiological recordings of AP waveforms were done at room temperature, and cells were held at –70 mV. We performed experiments at room temperature because spikes are shorter at 37 °C (0.7-0.8 ms)^17–19^, and are thus suboptimally sampled with our acquisition rate (440 Hz under 2P microscopy). Since GEVIs’ response time constants decrease with temperature at about the same rate as spike width, GEVI responses are often tested in vitro at room temperature^14,20–23^. We clamped GEVI-expressing cells to a range of typical AP waveforms and monitored changes in GEVI fluorescence. We used an AP waveform recorded from a representative hippocampal neuron and modified it to have an amplitude of 100 mV and a full width at half maximum (FWHM) of 4 ms, to mimic the shape of layer 2/3 cortical neurons at room temperature^17^. Cells were stimulated with 5 AP waveforms with a FWHM of 4 ms at 5 Hz and 10 AP waveforms with a FWHM of 2 ms at 100 Hz. In the same protocol, we also recorded GEVI response to a modified burst of APs recorded from adult mouse somatosensory cortex Layer 5 pyramidal neurons, designed to mimic APs on top of subthreshold depolarizations. The spike burst exhibited a subthreshold depolarization of ∼24 mV (from a baseline voltage of -70 mV) and APs with amplitudes of 60–90 mV (from a subthreshold voltage of -56 mV). The APs in the spike burst were 3–4 ms FWHM.

To characterize the sensors’ fluorescence vs voltage curves under 2P, cells and hardware set up were prepared as described above for voltage clamping at room temperature, except the power of 1050 nm was set to 25-42 mW. Cells were held at -70 mV for 4 seconds, followed by a series of 1-second voltage steps (90, 70, 50, 30, 20, 0, -20, -40, -60, -80, -100, and -120 mV), each separated by 1 second at -70 mV. To reduce the amount of light to which the cells were exposed, a 64 x 64-pixel galvo-galvo scanning image (14.1 µs dwell time, 1.24 µm pixel size) was recorded once every second for the duration of the patch-clamp protocol.

#### Optical responses to spike waveforms

To quantify fluorescent response to spike waveforms, the videos were analyzed using the same mathematical steps described in the “High-throughput screening data analysis” section. We corrected the red (GEVI) fluorescence trace for photobleaching by dividing each trace by the best-fit three-term exponential or by using a sliding window of 3 frames. Fitting of the fluorescent trace for photobleaching correction was performed outside of any voltage step. For each cell, the five 4-ms AP waveforms were averaged before calculating the peak response amplitude and spike width (FWHM).

#### Voltage response curves

Cells were analyzed using in-house-developed custom Python software. Individual cells were segmented in each field of view (FOV) using Cellpose 3^24^. To isolate the cell edge, a Laplacian filter was applied to the binarized Cellpose3 mask. Pixels were excluded if they became saturated in any frame during the experiment or if their baseline fluorescence was less than 1.5 times the median background signal. Background fluorescence was defined as the signal from regions of the image that did not contain cells. Due to the relatively small number of frames and their wide temporal spacing, photobleaching was approximated by fitting a linear regression to the fluorescence decay of frames acquired at a membrane potential of -70 mV.

Photobleaching was corrected by multiplying the fluorescence time series by the inverse of the fitted bleaching decay function. ΔF/F₀ was calculated using the background-corrected signal and the bleach-corrected signal, with F₀ defined as the average fluorescence intensity at −70 mV. We then fit the calculated ΔF/F₀ to the logistic sigmoid 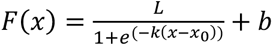 to determine the ΔF/F₀ at -70 mV. This value was used to offset all calculated ΔF/F₀ so that the fit function would have F(-70 mV) = 0 ΔF/F₀.

### AOD-based recordings in the mouse cortex using ULoVE

Animal experimentation was performed in accordance with the guidelines of the French National Ethics Committee for Sciences and Health, as outlined in the report on Ethical Principles for Animal Experimentation, in agreement with the European Community Directive 86/609/EEC, under agreement #29791.

#### Animal handling, viral injections and surgeries

5 male wild-type C57BL/6J adult mice (>P40, body weight 19–26 g) were housed under standard conditions (12-hour light/dark cycle, light on at 7 a.m., with water and food ad libitum). A preoperative analgesic was used (buprenorphine, 0.1 mg/kg), and Ketamine-Xylazine was used as an anesthetic (Centravet). The AAV2/1-EF1α-DIO-JEDI-2P-Kv-WPRE virus was produced in-house using a plasmid ratio (μg) of transgene:capsid:helper: 1:1.6:2. The AAV-EF1α-DIO-JEDI-2P-Kv-WPRE and AAV-EF1α-DIO-VADER1-TSER-Kv2.1 viruses were injected at a titer of 4x10^12^ GC/mL. To obtain a sparse density of GEVI-expressing neurons, AAV2/1 hSyn-Cre (Addgene, AV-1-PV2676) was co-injected at a final titer of 3.15x10^9^ GC/mL.

Viruses were diluted in a saline solution containing 0.001% pluronic acid (Thermo Fisher Scientific 24040032). 300 nL of each viral solution was injected at a flow rate of 100 nL/min into the cortex of each animal (coordinates from bregma: anteroposterior –2/–3.5 mm, mediolateral –2/–3 mm, and dorsoventral –0.32 mm from brain surface). Both indicators were expressed in the same animals, at different injection sites. A custom-designed aluminum headplate was fixed on the skull with layers of dental cement (Superbond). A 5-mm diameter #1 coverslip was placed on top of the visual cortex and secured with dental cement (Tetric Evoflow). Mice were allowed to recover for at least 15 days before recording sessions and co-housed (at least 2 mice per cage). Imaging was done between 19 and 64 days post-surgery. Before the awake recordings, mice were handled by the experimenter and progressively habituated to head fixation. Mice were handled before recording sessions to limit restraint-associated stress, and experiments were performed during the light phase of the cycle.

#### ULoVE voltage optical recording and experimental design

During recordings, mice were head-fixed via the metal cranial implant and placed on an unconstrained running wheel in the dark, where they were allowed to move spontaneously^25^. The duration of the recording sessions was limited to 3 hours. Recordings were performed using a custom-made random-access multiphoton system (Karthala System) equipped with acousto-optic deflectors (AODs)^26^. The excitation was provided by a femtosecond laser (InSight X3, Spectra Physics) mode-locked at 920 nm for JEDI-2P-Kv and 1045 nm for VADER1-tKv, with a repetition rate of 80 MHz. A water-immersion objective (CFO Apo25XC W1300, 1.1 NA, 2 mm working distance, Nikon) was used for excitation and epifluorescence light collection. The signal was passed through an IR-blocking filter (TF1, Thorlabs), split into two channels using a 562 nm dichroic mirror (Semrock), and directed to two H12056P-40 photomultiplier tubes (Hamamatsu) operating in photon-counting mode. A 510/84 filtered green channel was used to collect JEDI-2P-Kv signals, and a 607/70 filtered red channel was used for VADER1-tKv. Before the ULoVE recordings, JEDI-2P-Kv or VADER1-tKv-expressing neurons were located using the same AOD-based microscope, set to imaging mode. A time series of 50 images (1 μs/pixel, 0.091 μm/pixel) was acquired, and post hoc motion registration was performed to produce the images shown in the Figure 3A (left). Cell depth, indicated in Figure S3.1A, was measured from the surface of the brain. No selection criteria were followed for optical recordings.

Excitation was achieved using an optimized ULoVE pattern (Lombardini et al., in preparation) consisting of a series of 9 vertically aligned points (evenly spaced over a distance of 15 μm), multiplexed horizontally twice with a spacing of 2 μm. The 18 points are scanned diagonally (8 μm horizontally and 3 μm vertically) during the 50-μs acquisition time so that an extended excitation volume is homogeneously filled and the plasma membrane of the cell of interest can be continually recorded. For imaging, the laser power was set to deliver 15 mW post-objective and pre-sample, then adjusted to account for mono-exponential loss through tissue, with an attenuation length constant of 170 μm. For the 10-minute ULoVE optical recordings, the power used in the imaging mode was multiplied by 1.5 (i.e., 22.5 mW at the surface) to account for the greater excitation volume, while for the 60-minute recordings, the imaging power was multiplied by 1.2 (i.e., 18 mW at the surface). This smaller scaling factor reduced the photobleaching rate, allowing a longer recording period. The applied power never exceeded 200 mW at the brain surface. Using two patterns per cell provided a temporal resolution of 7142.9 Hz: 2 volumes x (50 µs acquisition time + 20 µs of AODs access time). The recordings shown in Figures 3A-F and S3.1 were stopped after 600 seconds (10 minutes), whereas those in Figures 3G-I and S3.2 were collected continuously for 60 minutes.

#### Signal analyses, spike extraction and waveform analyses

Each cell was recorded with two ULoVE volumes simultaneously, so the resulting trace represents the sum of photons collected from both volumes. The photon flux is the number of photons per second.

For photobleaching analysis, the raw trace was first fit with a bi-exponential function, and the resulting normalized fit was subtracted from the trace to correct for fast bleaching. This correction allowed further analyses of the full recording. The following slower bleaching phase (from 1 minute to the end) likely reflects a diffusion-related process and can be described by a power law. For this phase, the trace was low-pass filtered at 1Hz, down-sampled 140 times, and then linearly fit on log10-log10 scales, allowing us to extract the power law slope.

The normalized trace, corrected with the bi-exponential fit, was high-pass filtered at 0.5 Hz using a zero-phase distortion filter to remove slow drifts, and the % ΔF/F_0_ was calculated using the mean fluorescence signal as F_0_.

Spike detection was done using a custom-designed algorithm, as previously described. Briefly, raw voltage traces were first high-pass filtered with a second-order Butterworth filter at 40 Hz. Then we convert this trace into a delayed differential trace, which is the difference between the “signal” (defined as the trace’s mean over 2 ms) and the ’baseline’ (defined as the trace’s mean over 3 ms, delayed by 1 ms). The rationale is to define a baseline before the spike that excludes data points in the rising phase of the putative spikes. Choosing a baseline this way enables increasing contrast for both isolated spikes and spikes within bursts. A subsequent step consists of enhancing the contrast by multiplying the delayed differential trace by its positive high-frequency (250 Hz cutoff) content. Finally, this sharpened and highly contrasted trace is z-scored. Events above 20 standard deviations were considered as spikes. Lastly, spike onsets are extracted as the local peak of the linear regression slope over a 3 ms window on a 10 kHz linear interpolation trace.

For further spike quantifications, an average spike waveform was aligned on the spike onset. The spike amplitude was then taken as the peak value of the average spike waveform, measured from the onset. Full-width at half maximum (FWHM) is the extent in time at half maximum amplitude, taken from the average spike waveform after 20 kHz linear interpolation. Tauoff represents the time constant of an exponential fit on the repolarization phase of the spike. Dprim was also obtained from the average waveform, as previously published^27^. To calculate the spike SNR, the amplitude was divided by the shot noise, corresponding to the square root of the ΔF/F0 trace. For the 60-minute recordings, spike waveform metrics were obtained for periods of 300 s (5 minutes).

To evaluate the impact of downsampling on spike amplitude detection, we analyzed extracellular waveforms originally sampled at 7.1 kHz. Waveforms were upsampled to 1 MHz and then downsampled to rates between 25 and 7000 Hz. For each sampling rate, we tested 20 temporal phases by systematically shifting the downsampling starting point. Peak amplitudes were normalized to the original waveform maximum. We quantified amplitude loss and variability (coefficient of variation) across neurons and phases for both JEDI-2P and VADER1 recordings.

### Simultaneous electrophysiology and resonant-scanning two-photon voltage imaging in the mouse cortex

#### Animal handling

All procedures were carried out in accordance with the ethical guidelines of the National Institutes of Health and were approved by the Institutional Animal Care and Use Committee (IACUC) of Baylor College of Medicine. Mouse strains were sourced from our breeding colony. Mice were housed under standard conditions (12-h light/dark cycles, light on at 6 a.m., with water and food ad libitum).

#### Cranial windows, headbars, and viral injections

Cranial window surgeries were performed on C57BL/6J, NDNF-FlpO (RRID: IMSR_JAX:034876), Tlx3-Cre (RRID: MMRRC_041158-UCD), or SST-Cre mice (RRID: IMSR_JAX:013044) (7-16 weeks old). The NDNF-FlpO, Tlx3-Cre, and SST-Cre mice were used in experiments that did not require Flp or Cre recombinase due to the limited availability of wild-type mice. In total, seven male mice and five female mice were used in this study. Anesthesia was induced with 3% isoflurane and maintained between 1.5-2.0%. Meloxicam (5 mg/kg) was administered at the start of surgery for analgesia. Mice were placed in a stereotaxic frame (Kopf Instruments or Neurostar Drill and Injection Robot), with body temperature kept at 37℃ via a homeothermic blanket (Somnosuite with RightTemp Module, Kent Scientific). The scalp was shaved, and 0.05 cc of 0.5% bupivacaine (Marcaine) was subcutaneously injected into the incision site. After 10-20 minutes, approximately 1 cm² of skin was removed above the skull. The exposed fascia was cleared, and the wound edges were sealed with surgical adhesive (VetBond, 3M). A 13-mm stainless steel washer (Seastrom), attached to a custom, removable headbar, was implanted on the skull with dental cement (C&B Metabond). The washer was positioned about 2.7 mm lateral to the midline and 1.5 mm anterior to the lambdoid suture to provide optical access to the visual cortex. Once the cement had set, mice were freed from the stereotaxic head holder and secured on a small stage. A 4-mm diameter craniotomy was drilled at the center of the washer using a 1/2 mm burr (Meisinger HM1-005-HP). The exposed cortex was rinsed and submerged with artificial cerebrospinal fluid (ACSF; 125 mM NaCl, 5 mM KCl, 10 mM Glucose, 10 mM HEPES, 2 mM CaCl_2_, 2 mM MgSO_4_). After the craniotomy, the virus was injected into the visual cortex using a nano-injection pump (World Precision Instruments or Neurostar Drill and Injection Robot) at 5 nL/sec. The pipette was withdrawn approximately 1.5 minutes after the injection finished. During two-photon guided patching experiments, the dura was left intact to minimize swelling. A 4-mm glass coverslip (Warner Instruments) was sealed onto the skull with cyanoacrylate glue (VetBond), and the edges of the window were secured with super glue (Loctite). All mice were housed individually and given two weeks to recover post-surgery to allow virus expression.

VADER1-tKv expression was transduced by injecting either AAV-hSyn-VADER1-TSER-Kv2.1 (titer:7.22x10^12^ GC/mL) or AAV-hSyn-VADER1-TSER-Kv2.1 (titer: 1.3x10^13^ GC/mL) into the cortex. Mice were injected with total volumes ranging from 280-700 nL of virus at a depth of approximately 300-450 µm from the cortical surface.

After confirming viral expression, the initial cranial window was replaced with a custom coverslip that had a pre-drilled ∼0.5 mm hole (created using a diamond-tipped burr, Coltene/Whaledent). This replacement was performed at least 2 weeks post-surgery to allow for recovery and sufficient viral expression and to enable precise placement of the modified coverslip. The pre-drilled coverslip allowed patch pipettes to access the brain and was positioned to readily target VADER1-tKv expressing neurons. To prevent glue from leaking into the brain, the hole was sealed with a thin layer of Kwik-Cast (World Precision Instruments) or temporarily covered with ACSF or Gelfoam (Pfizer Hospital 00009-0353-01). Once the window was sealed, ACSF was applied to the window surface as an external solution to prevent the brain from drying out.

Traces shown in Figure 4B are from juxtacellular patch recordings. Throughout *in vivo* two-photon guided imaging experiments, mice were kept anesthetized using 1-2% isoflurane, and its body temperature was maintained at 37°C using a homeothermic blanket. *In vivo* two-photon-guided patching experiments are from 10 neurons acquired across six mice (four male, two female).

#### Electrophysiological recordings

Patch pipettes were pulled from borosilicate glass (Sutter Instrument; 1.5 mm outer diameter, 0.86 mm inner diameter) to a resistance of 6–11 MΩ and filled with ACSF to get juxtacellular recordings. Alexa Fluor 488 (20 μM) was added for pipette visualization. Pipettes were advanced under two-photon guidance using high positive pressure (∼400 mbar) to penetrate the dura and low pressure (20-40 mbar) when approaching target cells. Pressure was controlled via a manometer (Fisher Scientific 06-664-19) and a custom-built manifold that allowed rapid switching between high and low pressures, minimizing the ejection of internal solution. Recordings were acquired in current-clamp mode. Liquid junction potentials were not corrected. Patch-clamp recordings were performed using a Scientifica PatchStar micromanipulator and an Axon Instruments CV-7B headstage. Signals were amplified with a MultiClamp 700B amplifier (Axon CNS Molecular Devices) and digitized at 10 kHz using a National Instruments DAQ system (BNC-2090A interface) and recorded through a custom program written in LabVIEW 2016. A Quest Scientific HumBug noise eliminator was used in most recordings to suppress line noise.

#### Voltage imaging instrumentation

All scans were recorded using a ThorLabs two-photon resonant-scanning system with excitation provided by a tunable Ti:sapphire femtosecond laser (Coherent; Chameleon Vision). The frame rate for these experiments was either 440 or 720 Hz, and the excitation wavelength was 1030 nm. A 0.8 NA 16x objective (Nikon; N16XLWD-PF) was used for imaging. Output power was measured post-objective using a ThorLabs PM100D meter and ranged from 50 to 60 mW. Imaged cells were 60-170 μm from the brain surface. The imaging resolution ranged from 0.92 to 1.77 μm/pixel along the x-axis and 0.83-1.68 µm/pixel along the y-axis (Table S4.1).

#### Data exclusion criteria

Cells were excluded from the analysis if any of the following conditions were met: the spike-triggered average of the optical signal aligned to electrophysiological spikes did not show a clear fluorescence response, suggesting a potential mismatch between the patched and imaged cell; the fluorescence trace did not exhibit discernible spike-like deflections and was too noisy to reliably identify optocal spikes; tissue damage occurred due to high laser power; or substantial Z-axis motion occurred during imaging. In juxtacellular patch recordings, movement during pipette approach or tissue stabilization, or too much negative pressure can sometimes disrupt the membrane, allowing partial access to intracellular signals. These recordings were retained unless the cell died within the first 60 seconds; in that case, they were excluded. In addition, patch quality and cell health are expected to decline over time, potentially leading to patch loss or deterioration of the recording. To account for this, recordings were manually truncated to remove periods corresponding to patch loss or declining signal quality. Although recordings often lasted up to ∼10 minutes, analyses were restricted to the first minute to maximize signal quality and minimize the impact of photobleaching.

In a few recordings, the 60-second analysis window was selected later in the recording if there was no initial electrophysiological spiking or if the fluorescence signal was excessively noisy. After applying these exclusion criteria, ten patched cells remained.

#### Data pre-processing

Following each two-photon imaging experiment, the scans were corrected for motion as described in the voltage imaging data pre-processing section below. For each recording, we manually segmented cells within the FOV, then removed the dimmest 25% of pixels within the mask. Fluorescent traces were high-pass filtered at 10-Hz and z-score normalized by subtracting the mean and dividing by the standard deviation computed across the entire filtered trace. Electrophysiological traces were also 10-Hz high-pass filtered.

#### Optical spike inference

For Figure. 4B-D, 4N-O, spikes were detected in the optical trace using an adaptive threshold modeled after Caiman Volpy’s adaptive threshold^28^. An adaptive method was used to avoid manual, trace-specific threshold selection. Fluorescence traces were first high-pass filtered at 10 Hz using a fifth-order Butterworth filter. Then the traces were passed through the adaptive threshold function, which selects a threshold based on the statistical distribution of local peak amplitudes. After filtering, a first-pass peak detection was run to identify local maxima. To determine the detection threshold, the amplitude distribution of all candidate peaks was estimated using a Gaussian kernel density estimator (KDE). The noise distribution was approximated by reflecting the KDE around the median peak amplitude, assuming symmetric noise. Tail integrals of the signal (*F_max_*) and noise (*F_noise_*) distributions were computed across a range of amplitudes, and a score function:

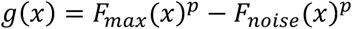

was evaluated, where *p ∈* (0,1] controls detection stringency (set to 0.1 in our analysis). The threshold was defined as the amplitude 𝑥 that maximized 𝑔(𝑥), and peaks exceeding this threshold were classified as optical spikes.

#### Precision-recall curves

Spikes were detected in both electrophysiological and optical traces. Fluorescence traces were high-pass filtered at 10 Hz using a Butterworth filter, then z-score normalized as previously described in “Data pre-processing”. Optical spikes were detected by applying the adaptive thresholding as described in “Optical spike inference”. To evaluate detection performance across a range of thresholds, the computed adaptive threshold was multiplied by a series of factors (typically 0.1-2.5), producing a set of binarized spike detection traces for each scale factor.

Detected electrophysiological spikes were mapped to the imaging frame immediately preceding their time. Spikes were grouped into events based on their temporal spacing: spikes separated by ≤ 18 ms were considered part of the same burst, and a refractory exclusion window of 29 ms was enforced before a new event could begin. Events were classified as singlets, doublets, and triplets+ based on the number of spikes within each event.

Detection performance was evaluated at the event level using a lenient matching criterion. An optical spike was considered a true positive (TP) if it occurred within the frame range spanning the start and end of a ground truth spike event, with a tolerance of ±1 imaging frame. Each optical spike could only be matched to one event. Ground-truth events without a corresponding optical detection were classified as false negatives (FN), and unmatched optical spikes were classified as false positives (FP). Precision, recall, and F1 score were then computed for each threshold scale factor:

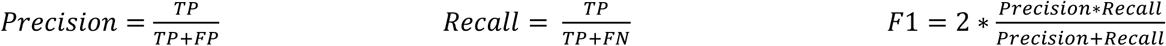

For each cell, precision-recall curves were constructed (Figure S4.2C) by plotting precision versus recall across all threshold scale factors. The best detection performance was defined as the threshold that maximized the F1 score for that cell (Figure 4D).

#### Spike-triggered average

Electrophysiological spikes were detected from patch-clamp recordings and aligned with imaging by assigning each spike to the imaging frame immediately preceding its time. Fluorescence traces were restricted to the analysis window, high-pass filtered at 10 Hz with a fifth-order Butterworth filter, and z-score normalized as described in the section “Data pre-processing”. Spike-triggered averages (STAs) were computed by extracting peri-spike samples from the z-scored trace around each spike-aligned frame (30 ms pre-spike, 30 ms post-spike) and averaging across events. SEM was computed across individual peri-spike snippets at each time point. Grand-average STA (Figure 4C) was computed by first calculating per-cell STAs and then interpolating each per-cell STA to a common 1-ms time grid before averaging across all six cells (mean ± SEM across cells). Per-cell STAs were computed for spikes occurring within the 60-second temporal window. The grand-average STA was computed across 6 cells imaged at 720 Hz.

#### Response amplitude analysis

For spike response amplitude quantification (Figure S4.2A-B), analyses were restricted to the defined 60-second region in selected cells. Fluorescence traces were high-pass filtered at 10 Hz using a fifth-order Butterworth filter and z-score normalized. Electrophysiological spike times were mapped to the nearest imaging frame to obtain normalized fluorescence values for individual singlet events.

### *In vivo* two-photon resonant imaging of VADER1-tKv responses in the visual cortex of mice

#### Animal handling & cranial window surgeries

As described in the previous section titled *Animal handling*, with the exception that dura was carefully removed while avoiding damage to improve imaging quality.

#### Voltage imaging instrumentation

Single-cell and multi-cell imaging scans shown in Figure 4E-4K, Figure S4.3-4.4, were acquired on a ThorLabs Bergamo resonant microscope with laser excitation provided by a tunable Ti:sapphire femtosecond laser (Coherent; Chameleon Discovery) tuned to 1050 nm or a two-photon resonant-scanning system (ThorLabs) with excitation provided by a tunable Ti:sapphire femtosecond laser (Coherent; Chameleon Vision) tuned to 1030 nm. Precompensation (7000 GDD) was applied internally in the laser via software. A dichroic mirror split the emission light into two channels: the green channel used a 525/50 nm filter, and the red channel used a 625/90 nm filter before being collected by two photomultiplier tubes. ScanImage (version 2018; Vidrio) was used to control the microscope and acquire imaging data. Imaging data on the Thorlabs Bergamo microscope was acquired with either a 16X 0.8 NA objective (Nikon; N16XLWD-PF) or a 25X 1.1 NA objective (CFI75 Apochromat 25XC W, Nikon Instruments).

#### Viral injections

VADER1-tKv shown in Figure 4F-4K, Figure S4.3-4.4 was transduced by injecting WT or Tlx-Cre with AAV-EF1α-DIO-VADER1-TSER-Kv2.1 (titer:2.00 x 10^13^ GC/mL). Cre-dependent VADER1 constructs injected in WT mice were co-injected with AAV1-CaMKII-0.4.Cre.SV40 (Addgene viral prep # 105558-AAV; 1.1 x 10^13^ GC/mL) in a 1:1-10 ratio. A total volume of 400-1000 nL was injected at a depth of approximately 300 and 500 µm from the cortical surface.

#### Sampling

Single-cell imaging data shown in Figure 4F-H, Figure S4.3, and Figure S4.4 were obtained from 6 neurons from 2 male and 1 female mouse, at depths ranging from 128 to 450 µm, with post-objective powers ranging from 70 to 90 mW, as measured using a ThorLabs PM100D meter. Imaging FOVs of the data shown in Figure 4E-H, Figure S4.3, and Figure S4.4 spanned 11-37 µm in width and 15-16 µm in height, with a resolution of 0.75-1.5 pixels/µm in x and 0.76-1.6 px/µm in y. Single-cell functional scans of spontaneous activity were acquired at 396 Hz. Functional imaging of responses to oriented visual stimuli shown in Figure 4J-K was obtained from a female mouse at a depth of 256 µm from the cortical surface. Post-objective power measured 50 mW using a ThorLabs PM100D meter. The imaging field-of-view spanned 83 × 111 µm in height and width, with a resolution of 0.9 px/µm, and was acquired at 198 Hz. Additional functional imaging scan of responses to visual stimuli not shown was acquired at a depth of 313 µm with 50 mW of post-objective power. FOV spanned 71 µm in width x 97 µm in height with resolutions of 0.3 px/µm in x and 0.4 px/µm in y at 396 Hz.

#### Oriented visual stimuli

Single-cell spontaneous activity scans were performed in the dark without any visual stimulation. Pixelwise orientation tuning scans utilized a global directional parametric stimulus (monet_voltage) as described at the end of this section. Custom MATLAB (version 2023b, MathWorks) and LabVIEW (version 2016, National Instruments) code was used to control the acquisition and synchronization of imaging and visual stimuli. For pixelwise-tuning scans, mice were shown Gaussian noise with coherent orientation and motion (monet_voltage). The stimulus contained 16 motion directions randomly mixed, each repeated 100 times. Each motion bout lasted 0.5 seconds. The total stimulus duration was approximately 14 minutes.

#### Data exclusion criteria

Scans with average x-y motion exceeding 10 μm, sudden artifacts, or loss of fluorescence due to water loss or external light contamination were excluded from analysis.

#### Data pre-processing

Voltage imaging data were pre-processed using a pipeline written in Python (3.6.9). Imaging data were motion-corrected in the horizontal plane to account for shifts in the x and y axes using a two-step algorithm. First, large motion shifts were corrected by creating a global average template across the entire scan and then calculating the phase correlation between this template and individual frames. Second, smaller motion shifts were addressed by generating a local template from averaged segments of different sizes (ranging from 3 to 30 seconds, depending on which yielded the best results) and computing the phase correlation between this local template and individual frames. After motion correction, individual somas were manually segmented from the motion-corrected summary image. Raw fluorescence traces were then extracted from the manually segmented masks based on the motion-corrected imaging data. Subsequently, extracted single-cell VADER1-tKv traces, shown in Figure 4F-K, Figure S4.3, and Figure S4.4, were high-pass filtered with a 10 Hz Butterworth filter and z-scored by subtracting the mean of the high-pass filtered trace before dividing the product by the standard deviation of the high-pass filtered trace.

#### Characterization of VADER1-tKv activity

To characterize the photostability of VADER1-tKv spiking shown (Figure 4F-H), spikes were detected by passing a threshold parameter to the find_peaks function using the scipy.signal Python package (version 1.2.3). Thresholds were defined as the sum of the mean of the trace and the standard deviation multiplied by 5. Singlets, shown in Figure 4G, were identified by finding isolated detected spikes that had at least 100 ms before and after another detected spiking event. Doublets, shown in Figure 4G, were identified by finding two detected spiking events within 15 ms of each other that had no events in the preceding 100 ms. The singlet spike-triggered average shown in Figure 4H is generated from detected singlets (n = 168) in the z-score trace shown in Figure 4F.

#### Pixel-wise analysis of orientation tuning

For the pixel-wise orientation-tuning analysis (Figure 4J-K), motion correction was applied, and the scans were normalized in two stages. First, the data were detrended by subtracting a 0.1 Hz low-pass-filtered version of the scans, then dividing by it. Second, the scans were normalized by subtracting their temporal means and dividing by the root-sum-of-squares of each pixel’s temporal data. A spatial median filter with a 3 × 3 kernel was applied to the scans to reduce noise. To determine each pixel’s preferred direction, a linear regression model was used. For each unique direction, a regressor *R(t)* was designed:

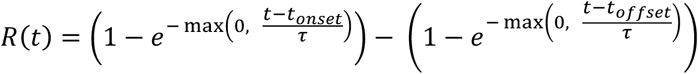

where *t* is the time of the frame, *t_onset_* is the start time of the trial, *t_o,set_* is the end time of the trial, and τ is a time constant, set to 0.7. The regression coefficients, indicating the response magnitude for each direction, were calculated by multiplying the detrended scan with the regressor matrix. For each pixel, a single complex number was obtained by projecting the N regression coefficients onto a complex-valued basis vector. This basis vector was created using *e2iθ*, where *θ* is the direction in radians, scaled by *1/√(N/2)*, where *N* is the number of unique directions. The orientation map was generated from the phase angle of this complex number, with each pixel representing the preferred orientation of motion that elicited the strongest response. The amplitude map was derived from the magnitudes of complex numbers, reflecting the strength of orientation tuning. The orientation map was then color-coded to show the preferred orientation of each pixel, while pixel brightness and saturation represented the tuning strength from the amplitude map. To generate orientation tuning curves, the mean change σ for each motion orientation was computed.

### *In vivo* two-photon resonant imaging of coexpressed GCaMP6s and VADER1-tKv activity in individual neurons in the visual cortex of mice

#### Animal handling & cranial window surgeries

As described in the previous section, “Animal handling”.

#### Voltage imaging instrumentation

Imaging data of single neurons coexpressing VADER1-tKv and GCaMP6s were acquired on the Thorlabs Bergamo microscope described in previous sections (see “Voltage imaging instrumentation”). Laser excitation was provided by a tunable Ti:sapphire femtosecond laser (Coherent; Chameleon Discovery) tuned to 1020 nm to acquire sufficient signal across both imaging channels. Imaging data were acquired with a 16×0.8 NA objective (Nikon; N16XLWD-PF) or a 25× 1.1 NA objective (CFI75 Apochromat 25XC W, Nikon Instruments).

#### Viral injections

To coexpress GCaMP6s and VADER1-tKv in the same cells, AAV-EF1α-DIO-VADER1-TSER-Kv2.1 (titer:2.00 x 10^13^ GC/mL) was injected mixed 1:1 with AAV1-CaMKII-0.4.Cre.SV40 (Addgene viral prep # 105558-AAV; 1.1 x 10^13^ GC/mL) or AAV8-hSyn-VADER1-tKv (titer 1.3 x 10^13^ GC/mL) mixed 1:1 with AAV1-hSyn-GCaMP6s (Addgeme viral prep #: 100843-AAV1, titer: 1.4 x 10^13^ GC/mL) into wild-type mice. Injection depths ranged from 250 to 400 µm from the cortical surface, with approximately 500 to 700 nL per injection site, with a maximum of two injection sites per cranial window.

#### Sampling

Single-cell VADER-GCaMP6s imaging data shown in Figure 4M-O were obtained from 6 neurons across 1 female and 2 male mice. Imaging depths ranged from 105 to 245 µm from the cortical surface, with imaging power ranging from 40 to 80 mW, as measured post-objective with a ThorLabs power meter (PM100D). Imaging FOVs spanned 24-50 μm in x and 24-32 μm in y. Imaging resolution spanned 0.75-1.5 um/px in x, and 0.76-1.6 μm/px in y. Imaging sampling rate was 396 Hz for all scans.

#### Data exclusion criteria

Similar exclusion criteria as previously described: scans with average x-y motion exceeding 10 μm, sudden artifacts, or loss of fluorescence due to water loss or external light contamination across both the green and red channels were excluded from analysis. Out of 32 total scans, 6 neurons from 3 mice were analyzed.

#### Data pre-processing

VADER1-tKv plus GCaMP6s imaging data were pre-processed through the same pipeline described in prior sections. Following motion correction, manual masks were segmented for both the green and red channels of the imaging field of view to extract raw fluorescence traces. The extracted single-cell VADER1 trace shown in Figure 4N and analyzed in Figure 4N-O was high-pass filtered using a 10 Hz Butterworth filter, and z-scored by subtracting the high-pass filtered trace from its mean, and dividing the product by the standard deviation of the high-pass filtered trace. The GCaMP6s signal shown in Figure 4N and analyzed in Figure 4N-O was low-pass filtered using a 20 Hz Hamming filter prior to z-scoring.

#### Characterization of simultaneous imaging data of VADER1-tkv-GCaMP6s activity in single neurons

To quantify the correspondence between VADER1-tKv spiking activity and GCaMP6s fluorescence shown in Figure 4O, we grouped VADER1-tKv spiking activity by the number of VADER spikes within a 50-ms window following an isolated VADER1-tKv spike, or two VADER1-tKv spikes that had no detected VADER1-tKv spikes within 500 milliseconds before the first spike’s onset. Detected spikes were determined as described in the previous section titled *Simultaneous electrophysiology and resonant-scanning two-photon voltage imaging in the mouse cortex – Optical Spike Inference* by using the same adaptive threshold function. Spike-triggered averages for the groups were computed on the 10 Hz Butterworth high-pass filtered ΔF/F_0_ VADER1-tKv trace and 20 Hz low-pass filtered ΔF/F_0_ GCaMP6s trace. We defined ΔF/F_0_ as (F-F_0_)/ F_0_, where F corresponds to the entire filtered trace, and F_0_ corresponds to the rolling mean of the bottom 10% of fluorescence values in a 60-second window of the entire filtered trace. Before averaging, each peri-event GCaMP6s was normalized to 0 by subtracting the mean of the trace sample from -150 to 0 ms preceding spike onset. Shaded areas are the standard error of the mean across all grouped, isolated single and double VADER1-tKv spike events.

### In vivo membrane localization assessment

#### Animal use & Mice preparations

All animal experiments were conducted in accordance with the National Institutes of Health guidelines for animal research. The Institutional Animal Care and Use Committee at the University of California, Berkeley approved procedures and protocols on mice.

All experimental mice were female, wild-type (C57BL/6J, stock no. 000664) purchased from Jackson Laboratories and bred in-house. Mice were 8 months old at the time of cranial window installation and were housed under a reverse light cycle.

Virus injection and cranial window implantation procedures have been described previouslyClick or tap here to enter text.^29^. Briefly, mice were anesthetized with isoflurane (1–2 % in oxygen) and given the analgesic buprenorphine (subcutaneously, 0.3 mg/kg). Animals were head-fixed in a stereotaxic apparatus (Model 1900; David Kopf Instruments). A 3.5 mm craniotomy was made over the intact dura covering the left visual cortex. In wild-type mice, the virus was injected to induce expression of genetically encoded voltage sensors using a glass pipette (Drummond Scientific Company) beveled to 35° and featuring a 15–20 μm tip opening, backfilled with mineral oil. A fitted plunger connected to a hydraulic manipulator (MO10; Narishige) was inserted into the pipette to load the virus and slowly inject it into 6–12 sites within the left visual cortex. At each injection site, 50 nL mixture of FSPAAV088-pAAV2/1-EF1a-DIO-VADER-A422V-TSER-kv2.1 WPRE (original titer 2 × 10^13^ GC/mL, diluted 6-fold in phosphate-buffered saline) and AV2/1-syn-Cre (original titer 3.15 × 10^13^ GC/mL, diluted 30-fold in phosphate-buffered saline) virus solution was injected at 250 µm below pia for voltage imaging. A glass window made of a single coverslip (No. 1.5; Fisher Scientific) was then embedded in the craniotomy and sealed in place using dental acrylic. Subsequently, a titanium head-post was affixed to the skull using cyanoacrylate glue and dental acrylic. *In vivo* imaging was conducted after ∼2 weeks of recovery and/or virus expression, and after habituation to head fixation. All imaging experiments were performed on head-fixed, awake mice 15 days post-surgery.

#### Microscope and imaging parameters

The ultrafast two-photon fluorescence microscope was described previously^30,31^. Briefly, light from a 1035 nm fiber laser (Monaco 1035-40-40, 1 MHz repetition rate; Coherent) was focused by a cylindrical lens and directed onto a pair of mirrors with a small tilt angle. Through free-space angular-chirp-enhanced delay (FACED), each laser pulse was split into multiple pulses that were spatially separated and temporarily delayed, forming a 1D array of excitation foci along X axis at the objective focal plane. The FACED foci array was scanned by a galvo mirror along the Y axis to generate one FACED FOV of 80 µm × 400 µm using a 25× 1.05 NA objective (XLPLN25XWMP2; Olympus). To extend the FOV, multiple FACED FOVs were tiled, yielding a 320-480 µm × 400 µm region.

The tiled FACED FOVs were post-processed with Y-axis shifts to better align structures and X-axis stitching to remove overlapping regions. The final FOV size was reported for each figure. The post-objective excitation power was 164 mW. Imaging was performed in the left visual cortex of the mouse, at depths ranging from 95 µm to 128 µm.

#### Image presentation in figures

Data were analyzed using custom MATLAB (Mathworks) programs. Time-streaming images were motion corrected using an iterative cross-correlation method^29^, the images presented in figures were illumination corrected along the FACED line scan to account for nonuniform excitation. Each image had its rows along the FACED line-scan direction and its columns along the Y galvo-scanning direction. Illumination correction was performed by first computing the average illumination profile along the FACED line-scan direction and then dividing each image row by this profile. This normalization corrected for nonuniform illumination across the FACED line scan while preserving relative signal variations.

## Data availability

The sequence of VADER1 is available from GenBank (accession number to be provided before publication). Plasmids used for AAV packaging for *in vivo* voltage imaging are available from Addgene (Addgene IDs to be provided before publication). Raw data for main and supplementary figures have been uploaded to Zenodo (to be provided prior to publication). Data that is too large to upload will be made available upon request.

## Code availability

This paper does not report the original code needed to analyze the data generated by this study. Any additional information required to reanalyze the data reported in this paper is available from the lead contact upon request.

## Acknowledgments

We thank Kim Tolias’ lab (BCM) for assistance with neuronal transfection protocols and for providing neurons for testing, Jun Chu (SIAT) for providing RFP/OFP constructs, and Nathan Shanner (UCSD) for assistance in RFP optimizations. We thank Fatima Nguyen (BCM) and Na Zhou (BCM) for help with stereotaxic surgeries. We thank Annick Ayon (ENS) and Bertrand Ducos (ENS) for virus titration. We thank Imane Bendifallah for cell culture preparation and transfection; and both Bendifallah and Ruth Sims (Vision Institute) for in vitro evaluation of a VADER1 intermediate.

## Grant support

The project described was supported in part by the Integrated Microscopy Core (supported by NIH grants P30 DK056338, P30 CA125123, and S10OD030414). The project was supported by a Vivian L. Smith Endowed Professorship in Neuroscience (FSP); a Klingenstein-Simons Fellowship Award in Neuroscience (FSP); the McNair Medical Foundation (FSP); Welch Foundation grant Q-2016-20190330 and Q-2016-20220331 (FSP); NIH grants R01EB027145 (FSP), R01EB032854 (FSP), U01NS113294 (FSP, ML), U01NS118288 (FSP), U01NS133971 (FSP, JR, LB), R01NS136027 (FSP, JR, VE), R01NS146023 (FSP, JR), R01NS146078 (FSP, JR), U01NS118300 (NJ), U01NS137449 (NJ), RF1NS128901 (JR), R34NS132045 (JR) ; Weill Neurohub (NJ); NSF grants NeuroNex 1707359 (FSP) and IdeasLab 1935265 (FSP); Swiss National Science Foundation grant Sinergia CRSII5_216632 (VE), a fellowship from Fondation des Neurosciences de Paris (TM); and Agence Nationale pour la Recherche grants ANR-25-CE16-7612-01 (LB) and ANR-20-CE16-0018-01 (VV).

## Author contribution

### St-Pierre lab

FSP conceived the project. FSP, SY, AJM, and XL coordinated the project. SY, AJM and XL screened GEVIs. SY, TT, AJM and XL benchmarked GEVIs in vitro. GF, ZL, XL and SY developed code to analyze screening or characterization data. XD, MS, SJ made GEVI stable cell lines. XD, SJ, SY and AJM constructed mutagenesis libraries and viral vectors. FSP, SY, AJM and XL prepared figures and wrote the manuscript.

### Bourdieu lab

VV, JB, and LB coordinated the project and wrote the manuscript. JB generated AAVs. V.V. performed mouse surgeries, conducted experiments and analyses. DGS wrote text, captions and perform figures production.

### Reimer lab

JR coordinated the experiments, analysis, and writing. MG and NH performed the VADER1 surgeries, imaging experiments, data analysis, and contributed to the writing. LR performed the VADER1/GCaMP6s surgeries and imaging. MH contributed to the data analysis pipelines. YY performed the pixelwise tuning.

## Competing interests

F.S.-P. holds a US patent for a voltage sensor design (patent #US9606100 B2) that encompasses the GEVI reported here. The remaining authors declare that they have no competing interests.

## Declaration of generative AI and AI-assisted technologies in the manuscript preparation process

During the preparation of this work, the author(s) used CoPilot in order to improve sentence-level structural refinement. After using this tool/service, the author(s) reviewed and edited the content as needed and take(s) full responsibility for the content of the published article.

