## Supplementary for "A red-emitting, genetically encoded indicator for two-photon voltage recording in vivo"

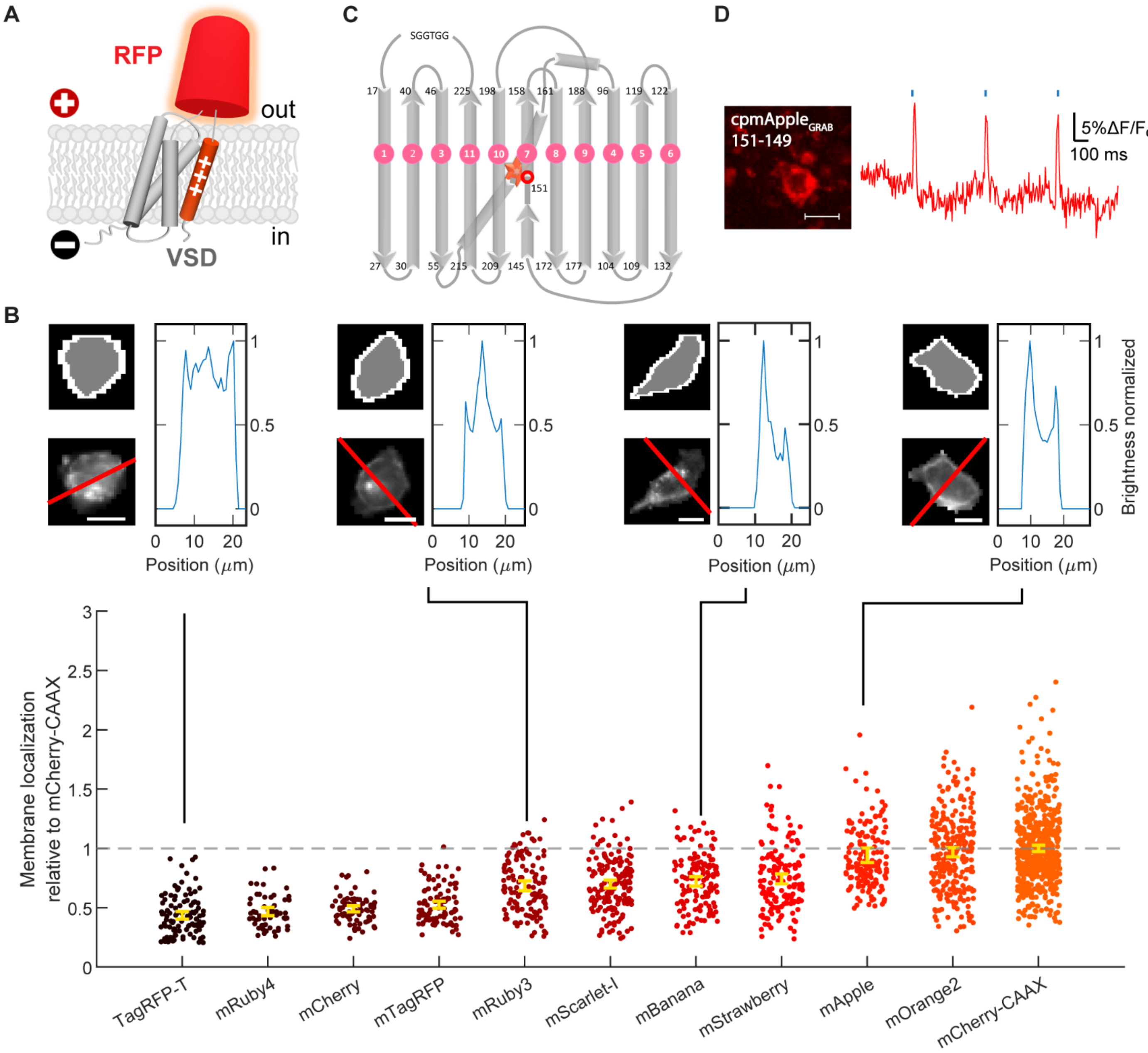


**Figure S1.1. Systematic identification of orange and red fluorescent proteins suitable for the replacement of circularly permuted GFP in the GEVI JEDI-2P.** (**A**) Conceptual schematic of membrane-localization screening constructs used to identify orange and red fluorescent proteins (OFPs and RFPs) for engineering red-shifted GEVIs. To assess fusion compatibility and extracellular folding independently of voltage-sensing function, native (non–circularly permuted) OFPs and RFPs were inserted into the extracellular loop between the third and fourth transmembrane helices of the JEDI-2P voltage-sensing domain (VSP), replacing the circularly permuted GFP. (**B**) Quantification of plasma membrane localization for the constructs described in (A). Membrane localization was measured as the ratio of the mean fluorescence intensity at the plasma membrane to the mean fluorescence intensity in the cytoplasm. Values were normalized to those obtained for membrane-anchored mCherry fused to a farnesylation sequence (CAAX). Due to spectral overlap, mCherry-CAAX was expressed in separate cells. Circles represent individual data points from n > 57 HEK293-Kir2.1 cells per construct. Yellow bars: means. Error bars: 95% CI. *Top*: Representative analysis examples for selected cells. In each group, the top-left panels show segmentation masks identifying plasma membrane (white) and cytoplasm (gray) pixels. The upper-right panels display fluorescence intensity profiles along the intersecting lines shown in red in the lower-left panels. Scale bar, 10 μm. (**C**) β-strand architecture of circularly permuted mApple, with strands numbered in pink, residue positions indicated at strand termini, and the chromophore indicated with a red star. The original N- and C-termini are connected by a short linker (SGGTGG). The red circle indicates the approximate location of the site around which circular permutation was introduced. (**D**) Membrane localization and voltage response of an early GEVI variant incorporating cpmApple. *Left*: Representative image of an HEK293-Kir2.1 cell expressing a GEVI incorporating a circularly permuted mApple with new N and C termini at 151 and 149, respectively. *Right*: Fluorescence responses of the same construct to electrical field stimulation (blue vertical lines). Traces show the mean fluorescence signal from an example field of view. Scale bar: 20 μm.

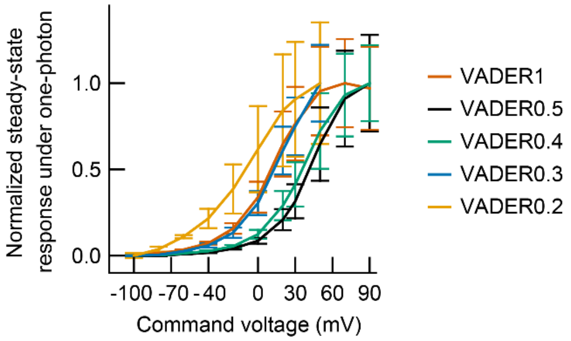


**Figure S1.2. Normalized voltage-response curves of VADER1 and screening intermediates under widefield one-photon illumination.** Voltage–response curves shown in Fig. 2G were normalized to their respective maximal fluorescence responses. n = 5 (VADER0.2), 11 (VADER0.3), 6 (VADER0.4), 5 (VADER0.5), 5 (VADER1) HEK293A cells. Error bars: 95% CI.

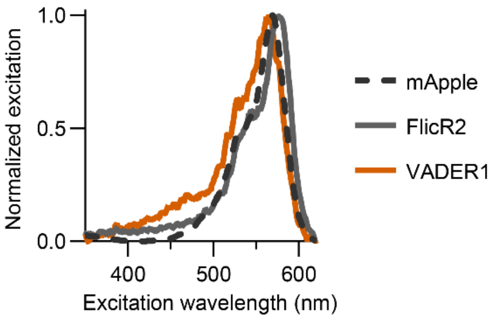


**Figure S2.1. VADER1's one-photon excitation spectra.** For each spectrum, values were normalized to the peak intensity. To recapitulate the membrane localization characteristic of GEVIs and thereby equalize experimental conditions, mApple was targeted to the plasma membrane using a prenylation (CAAX) sequence. Emission was collected at 650/10 nm. n = 6 (VADER1), 6 (FlicR2), 3 (mApple) independent transfections in HEK293A cells.

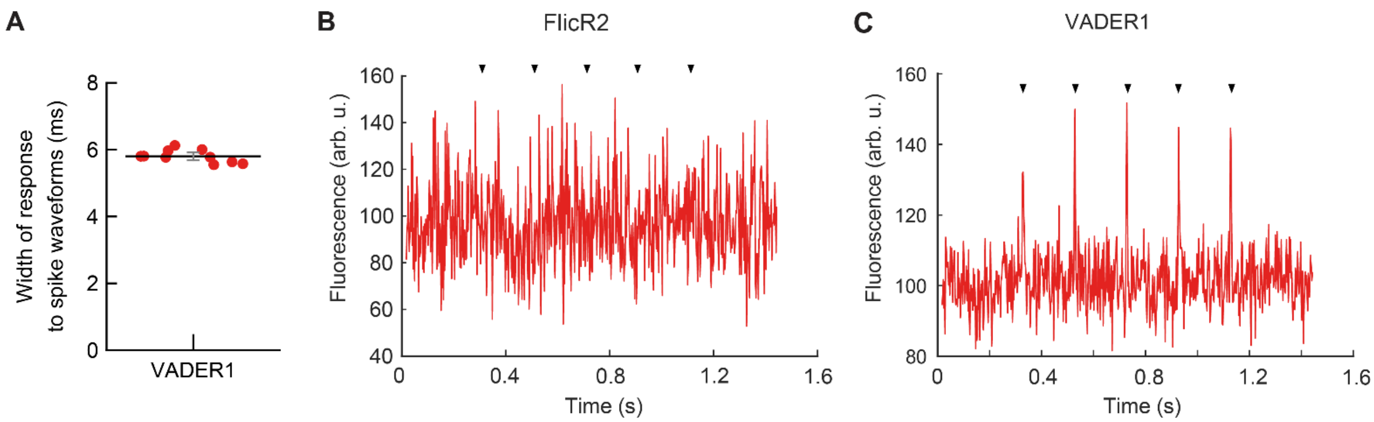


**Figure S2.2. Optical responses of VADER1 and FlicR2 to isolated spike waveforms under 2P resonant-scan imaging *in vitro*.** (**A**) Full-width-at-half maximum (FWHM) quantification of VADER1 fluorescence responses to isolated spikes (100-mV amplitude, 4-ms FWHM waveform). (**B**–**C**) Representative background-corrected, detrended fluorescence trace of FlicR2 (**B**) and VADER1 (**C**) acquired under two-photon resonant scanning in response to the same isolated spike waveform. Black triangles indicate the timing of the voltage spikes.

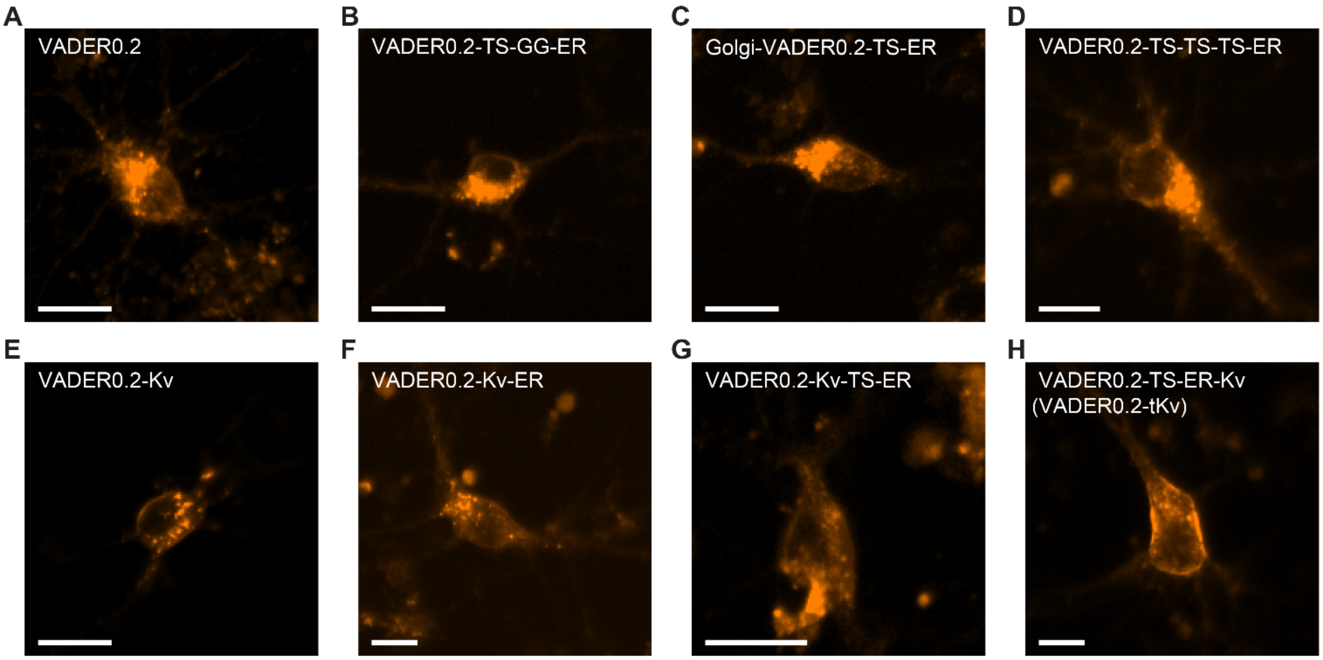


**Figure S2.3. Membrane localization of a VADER variant with different trafficking tags.** (**A-H**) Representative images of dissociated rat cortical neurons expressing VADER0.2 fused to individual or combinations of trafficking motifs. Constructs were expressed using a hSyn1 promoter. Trafficking motifs include: TS, the trafficking signal from Kir2.1[1]; ER, the endoplasmic reticulum export signal FCYENEV[1]; Golgi, the Golgi-export motif from Kir2.1[2], and Kv: the soma-restriction motif from Kv2.1[3]. GG denotes two consecutive glycines residues. For simplicity, the TS-ER-Kv combination is henceforth referred to as tKv. Images were acquired at DIV 14-15, 4-7 days after transfection. Scale bars: 20 μm.

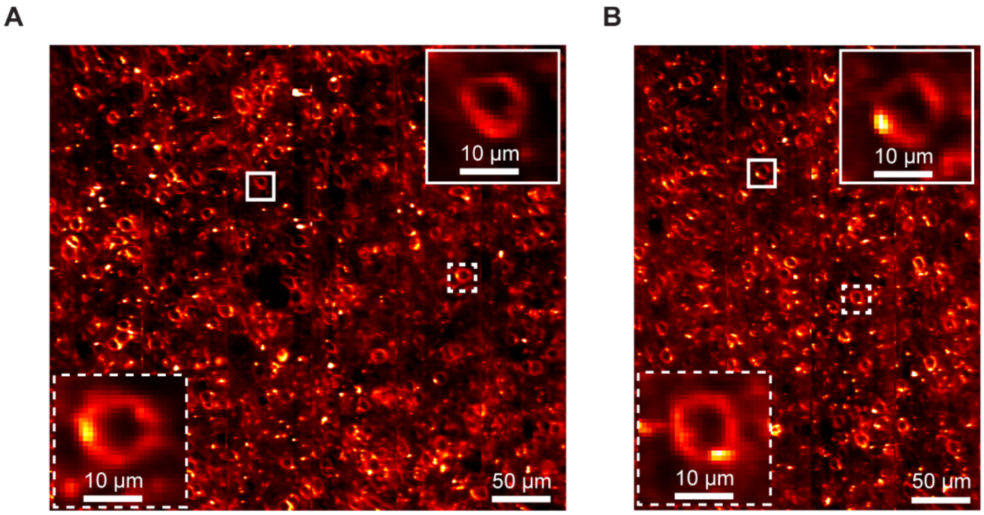


**Figure S2.4. Additional images illustrating the cellular localization of VADER1-tKv in the mouse cortex.** Fields-of-view (FOV) are located in layer 2/3 of visual cortex area V1. Images were acquired by two-photon scanning microscopy in a head-fixed mouse. Excitation: 1035-nm. FOV sizes are (**A**) 480 x 400 µm; (**B**) 320 x 400 µm. Both FOVs were obtained from the same animal.

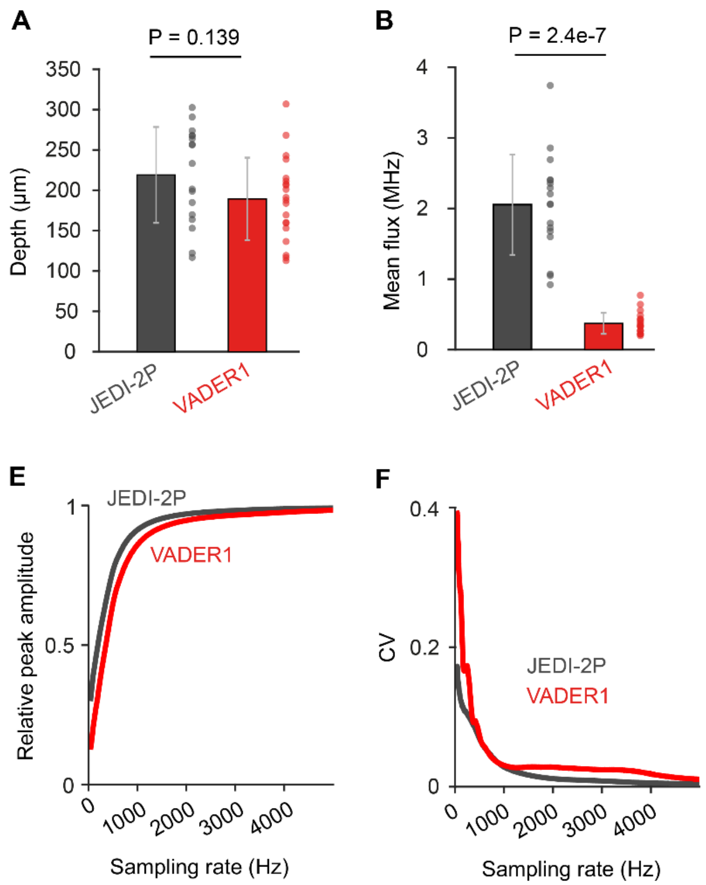


**Figure S3.1. Additional quantitative comparisons between JEDI-2P-kv and VADER1-tKv in vivo recordings, in awake mice.** (**A**) Depth, in µm, (**B**) mean photon flux, (**C**) spike Signal-to-Noise Ratio and (**D**) discriminability index of each of the 10-minute ULoVE recordings represented in Fig. 5A-F (VADER1-tKv: 20 cells, JEDI-2P-Kv: 17 cells). Two-sample Wilcoxon rank sum test; error bars represent the SD. (**E**) Relative peak spike amplitude (mean over 20 phases) of the 17 JEDI-2P and the 20 VADER1 expressing neurons plotted against sampling rate (down-sampled from their 1 MHz up-sampled average spike waveforms). (**F**) Coefficient of variation (CV), corresponding to the SD over the mean, plotted against sampling rate.

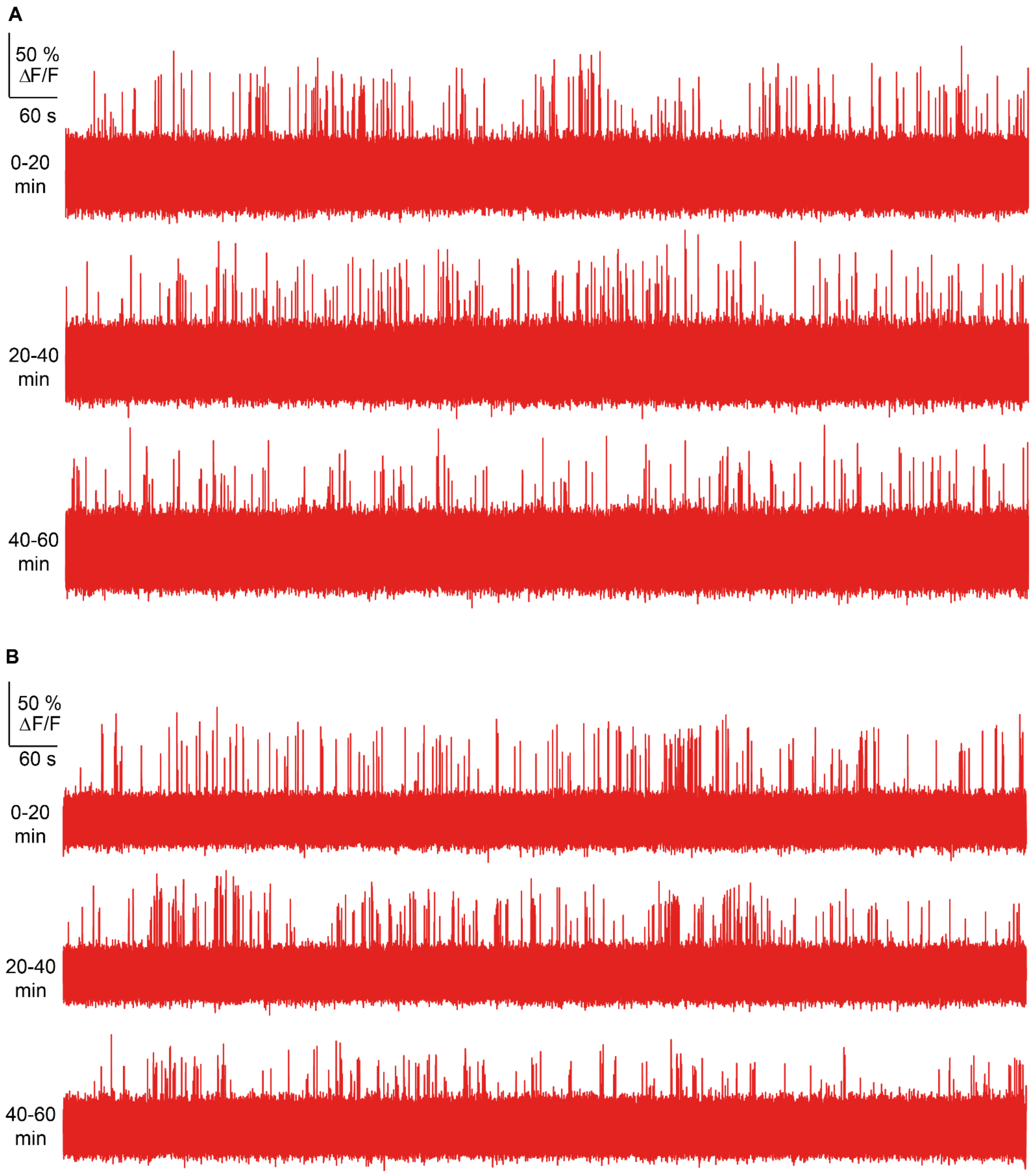


**Figure S3.2. Additional examples of 1-hr ULoVE recordings from neurons expressing VADER1-tKv**. (**A**) and (**B**) correspond to two additional layer 2/3 cortical neurons. Experimental conditions matched Fig. 3G, including the excitation wavelength (1045 nm), power at the brain surface (18 mW), and the acquisition sampling rate (7.2 kHz). The traces were smoothed with a Gaussian filter (time width: 0.4 ms).

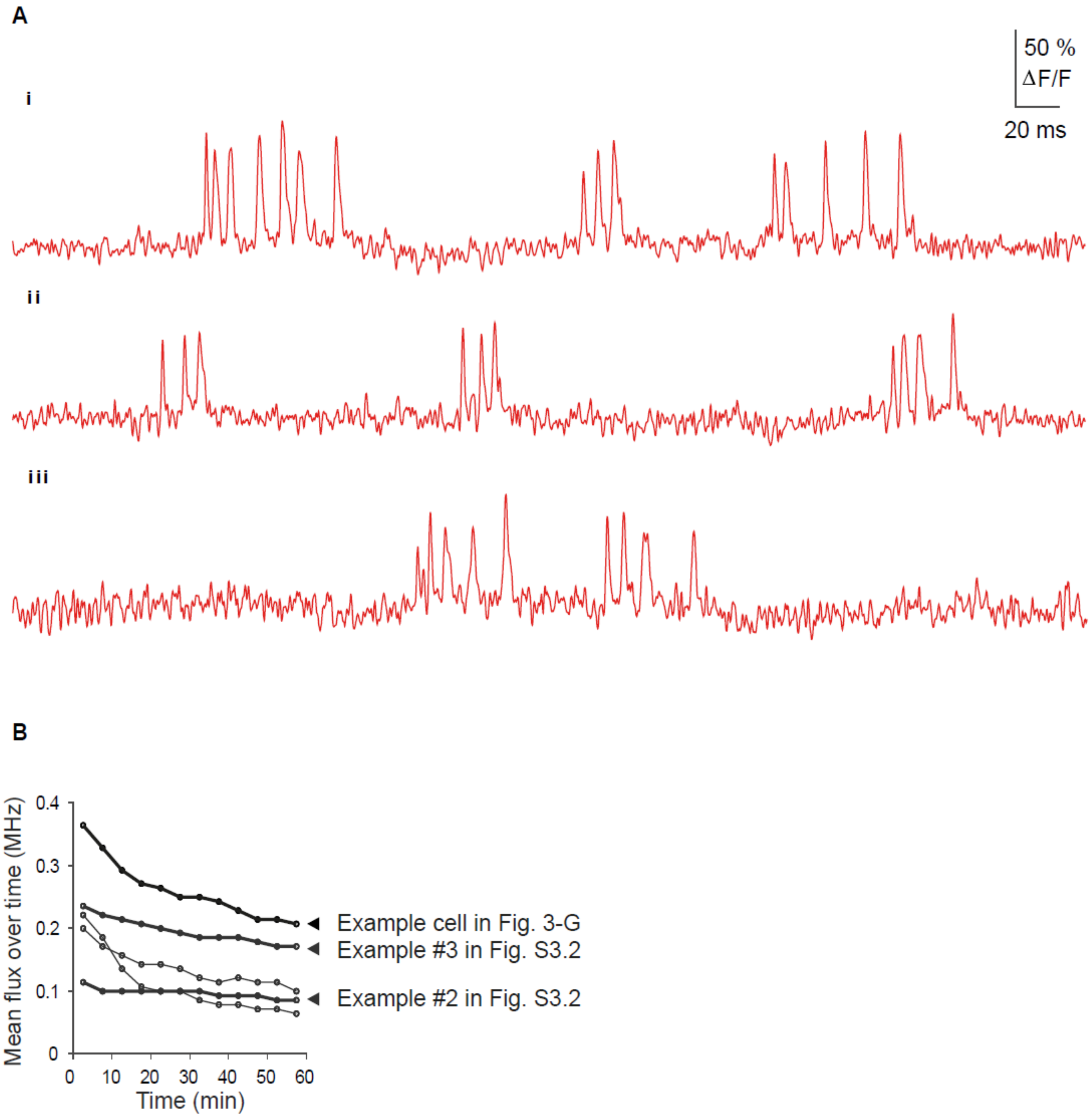


**Figure S3.3. Expanded views and photon-flux quantification from 1-hr ULoVE recordings of VADER1-tKv-expressing neurons**. (**A**) Expanded views of the recordings show in Fig. 3G (subpanels i-iii). (**B**) Quantification of changes in photon flux over 60 min for five L2/3 VADER1-tKv-expressing neurons. The dataset corresponds to that shown in Fig. 3H-I. Bold black traces indicate the example neurons in Fig. 3G and S3.2.

| Cell ID | Animal ID | # rows per FOV | # columns per FOV | FOV height (μm) | FOV width (μm) | Row height (μm) | Column width (μm) | # rows in cell mask | Laser flyback time (ms) | FPS | Zoom | Excitation power (mW) | Depth (μm) | Spike count (60 s) | Singlet count (60 s) |
|---|---|---|---|---|---|---|---|---|---|---|---|---|---|---|---|
| **1** | 1 | 18 | 64 | 30.2 | 113.3 | 1.68 | 1.77 | 13 | 0.2 | 720 | 11 | 55 | 83 | 102 | 56 |
| **2** | 2 | 30 | 64 | 25.0 | 64.6 | 0.83 | 1.01 | 20 | 0.3 | 440 | 21 | 38 | 104 | 10 | 10 |
| **3** | 2 | 30 | 64 | 25.0 | 64.6 | 0.83 | 1.01 | 17 | 0.3 | 440 | 21 | 45 | 168 | 20 | 18 |
| **4** | 2 | 30 | 64 | 22.8 | 59.0 | 0.76 | 0.92 | 19 | 0.3 | 440 | 23 | 45 | 165 | 57 | 55 |
| **5** | 3 | 18 | 60 | 19.7 | 56.6 | 1.09 | 0.94 | 14 | 0.2 | 720 | 16 | 50 | 127 | 222 | 190 |
| **6** | 4 | 16 | 64 | 24.6 | 103.8 | 1.54 | 1.62 | 13 | 0.3 | 720 | 12 | 43 | 144 | 108 | 97 |
| **7** | 4 | 16 | 70 | 18.5 | 69.2 | 1.15 | 0.99 | 12 | 0.3 | 720 | 13 | 42 | 92 | 35 | 18 |
| **8** | 5 | 18 | 64 | 17.5 | 75.3 | 0.97 | 1.18 | 11 | 0.2 | 720 | 18 | 67 | 130 | 69 | 67 |
| **9** | 1 | 18 | 64 | 21.1 | 64.3 | 1.17 | 1.00 | 13 | 0.2 | 720 | 14 | 65 | 69 | 79 | 27 |
| **10** | 6 | 30 | 64 | 25.0 | 64.6 | 0.83 | 1.01 | 20 | 0.3 | 440 | 21 | 74 | 133 | 72 | 36 |

**Table S4.1. Recording parameters for simultaneous two-photon imaging and juxtacellular patch-clamping.** Imaging was performed using resonant scanning. FPS, frames per second. Excitation power was measured at the objective. Imaging depth was calculated from the surface of the brain. Spike counts were measured within the 60-s epochs used for precision-recall analyses (Figure 4K, S4.4C). Singlet spikes were defined as events with no other detected spikes occurring within 18 ms before or 29 ms after the spike time.

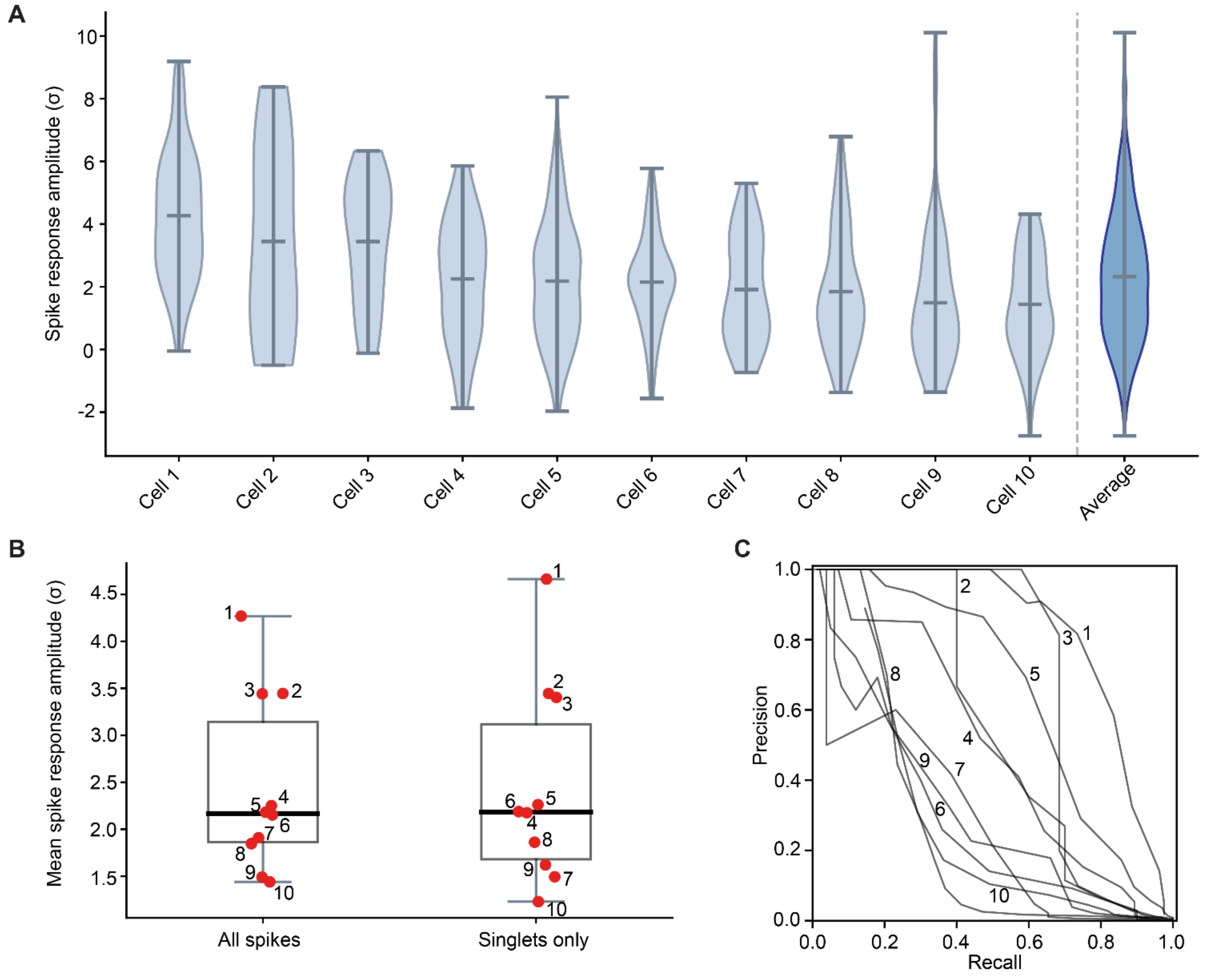


**Figure S4.2. Optical spike response distributions and spike-detection performance in individual cells.** (**A**) Distributions of z-scored optical responses to spikes for each cell. Center horizontal bars: mean responses. Top and bottom horizon bars indicate the maximal and minimal values, respectively. The rightmost violin plot shows the mean distribution across the 10 cells. (**B**) Mean optical spike amplitude per cell calculated from all detected spikes (left) or restricted to isolated singlet spikes (right). Boxplots indicate the interquartile range, with the median shown by the horizontal line. Individual cells are displayed as red points and labeled by cell number. (**C**) Precision–recall curves for optical spike detection across the 10 cells.

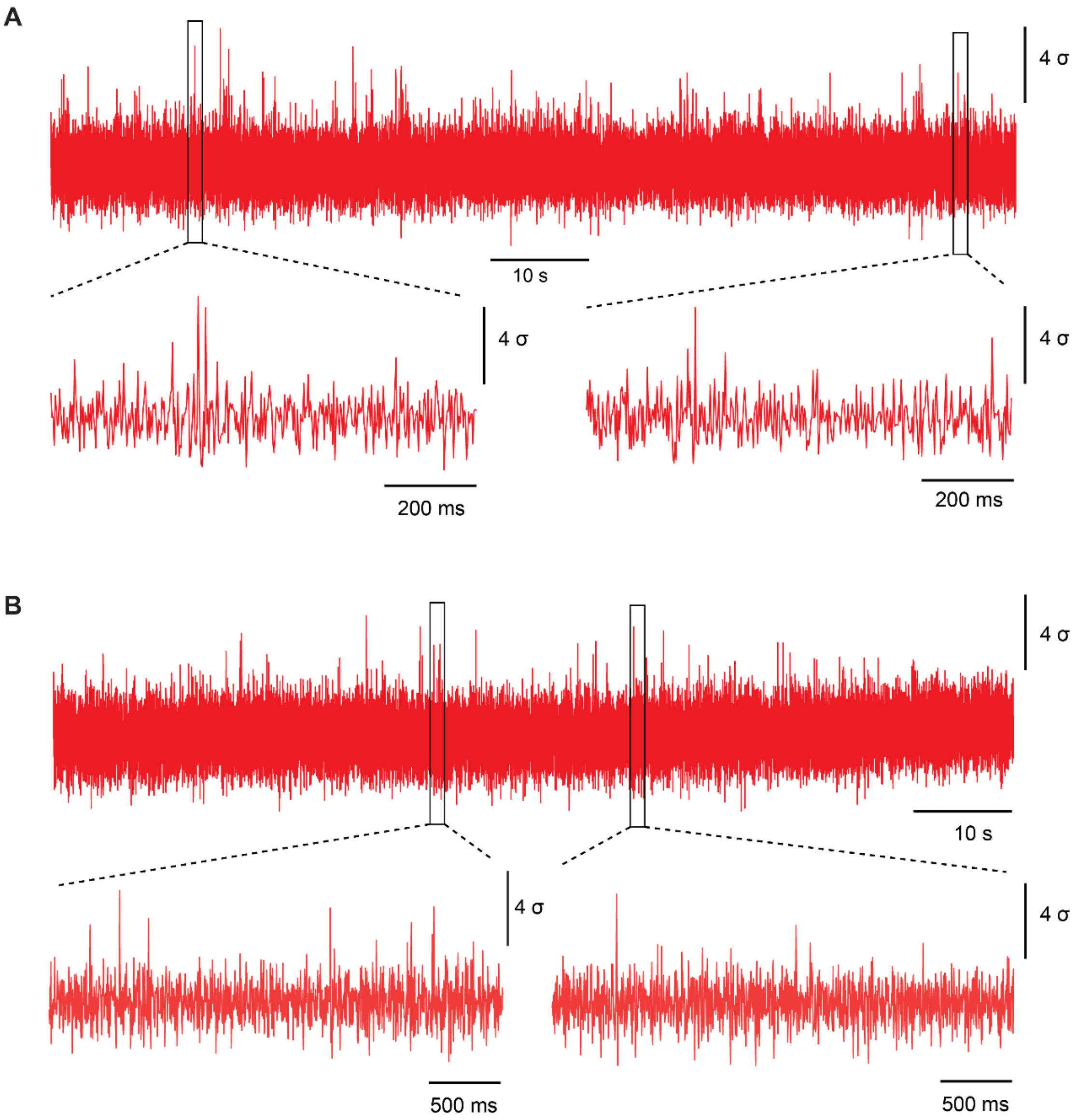


**Figure S4.3. Optical recording of spikes in Layer-2/3 pyramidal neurons using two-photon resonant-scan imaging of VADER1-tKv**. (**A-B**) Recordings from two individual neurons. Expanded views of spiking epochs near the beginning and near the end of each recording are shown below the corresponding full traces. Expanded views of spiking epochs near the start and the end of the recordings are shown below. Acquisition framerate: 396 Hz. Excitation wavelengths: 1030 nm. Both neurons are from the same mouse.

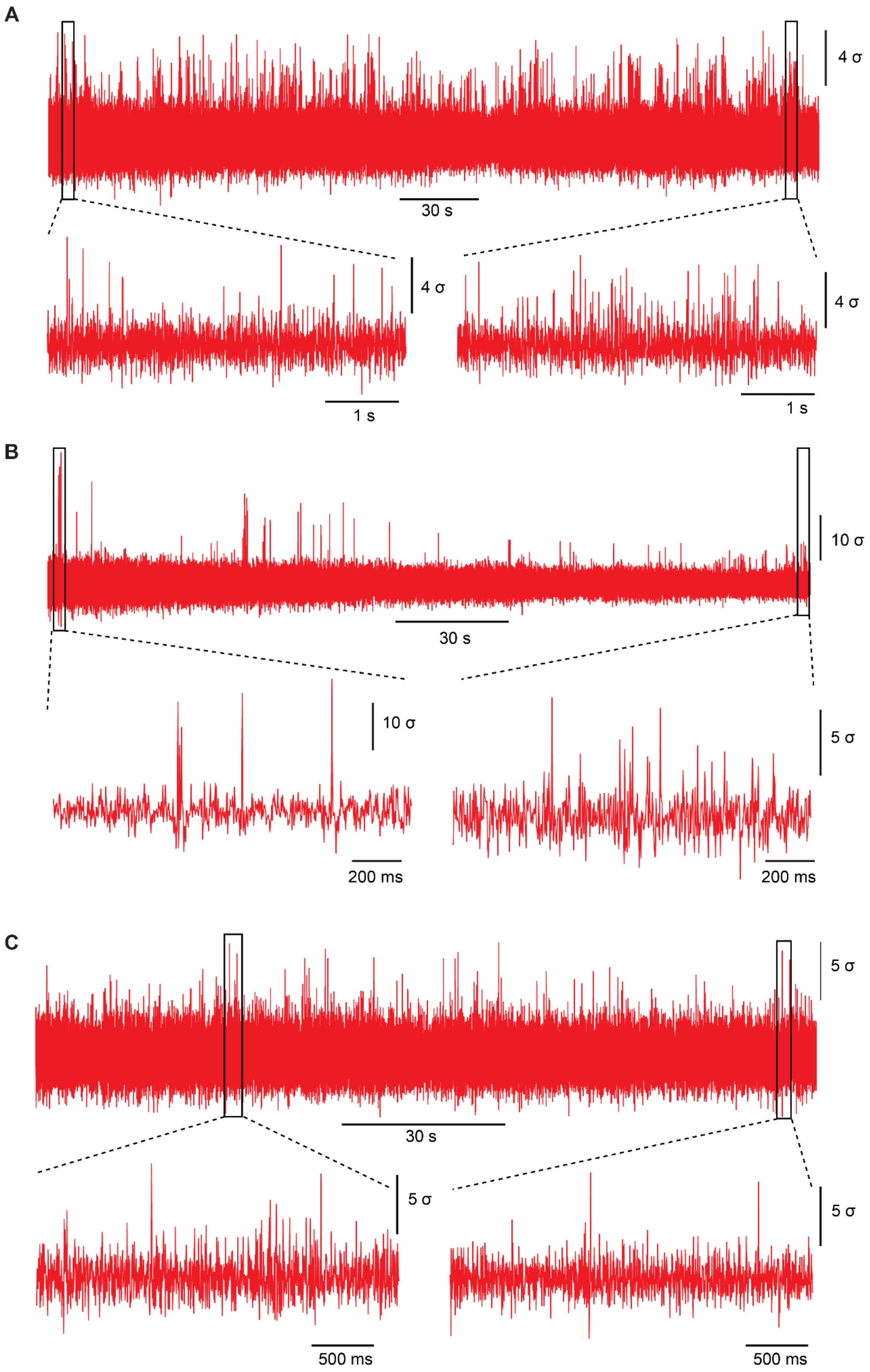


**Figure S4.4. Optical recording of spikes in Layer-5 pyramidal neurons using two-photon resonant-scan imaging of VADER1-tKv**. (**A-C**) Recordings from three individual neurons. Expanded views of spiking epochs near the beginning and near the end of each recording are shown below the corresponding full traces. Expanded views of spiking epochs near the start and the end of the recordings are shown below. Acquisition framerate: 396 Hz. Excitation wavelengths: 1050 nm (A), 1030 nm (B,C). Neurons here and from Fig. 4H are from 2 mice.
